# The impact of ambient contamination on demultiplexing methods for single-nucleus multiome experiments

**DOI:** 10.1101/2025.02.06.636969

**Authors:** Terence Li, Marcus Alvarez, Cuining Liu, Kevin Abuhanna, Yu Sun, Jason Ernst, Kathrin Plath, Brunilda Balliu, Chongyuan Luo, Noah Zaitlen

## Abstract

Sample multiplexing has become an increasingly common design choice in droplet-based single-nucleus multi-omic sequencing experiments to reduce costs and remove technical variation. Genotype-based demultiplexing is one popular class of methods that was originally developed for single-cell RNA-seq, but has not been rigorously benchmarked in other assays, such as snATAC-seq and joint snRNA/snATAC assays, especially in the context of variable ambient RNA/DNA contamination. To address this, we develop ambisim, a genotype-aware read-level simulator that can flexibly control ambient molecule proportions and generate realistic joint snRNA/snATAC data. We use ambisim to evaluate demultiplexing methods across several important parameters: doublet rate, number of multiplexed donors, and coverage levels. Our simulations reveal that methods are variably impacted by ambient contamination in both modalities. We then applied the demultiplexing methods to two joint snRNA/snATAC datasets and found highly variable concordance between methods in both modalities. Finally, we develop a new metric, *variant consistency*, which we show is correlated with cell-level ambient molecule fractions in singlets. Applying our metric to two multiplexed joint snRNA/snATAC datasets reveals variable ambient contamination across experiments and modalities. We conclude that improved modelling of ambient material in demultiplexing algorithms will increase both sensitivity and specificity.

## Introduction

Single-cell sequencing has enabled scientists to study biological systems at fine-scale resolution^1,2,3,4^. Recently, significant effort has been devoted to understanding the impact of genetic variation in single-cell RNA sequencing (scRNA-seq) experiments, which requires large sample sizes^5,6,7,8,9,10,11^. To achieve this scale and reduce technical variation, researchers use multiplexed designs in which multiple samples are processed simultaneously. Genotype-based demultiplexing methods are then used to infer cell identities by leveraging genetic differences between donors. In addition, genotype-based demultiplexing methods are capable of detecting between-donor (heterotypic) doublets, which cannot be accurately identified by traditional feature-based doublet callers^12,13,14^. To date, many genotype-based demultiplexing methods^15,16,17,18,19,20,37,38^ have been developed explicitly for scRNA-seq and have been shown to be accurate in scRNA-seq data^21^. More recently, they have also been successfully applied to joint single nucleus RNA/ATAC sequencing experiments^23^.

One question that has not been rigorously explored is the extent to which ambient RNA/DNA impacts demultiplexing method accuracy in these assays. Ambient RNA/DNA contamination arises from the capture of cytoplasmic molecular material and barcode swapping events^22^ and is an especially prevalent issue in single nucleus assays^27^. Such contamination can impact demultiplexing performance because genetic variants from other multiple donors will be present in contaminated droplets **(Fig 1A)**. To date, multiple studies have examined demultiplexing method performance in both scRNA-seq and scATAC-seq^21,23,24,25,40,41,42,43^ and shown that methods tend to identify similar numbers of singlets both within and across modalities. However, these studies benchmarked demultiplexing across low ambient RNA rates (5 - 10%) that may not be reflective of single-nucleus data^27,39^ and have not explored the effects of ambient DNA on snATAC-seq data.

**Figure 1:**
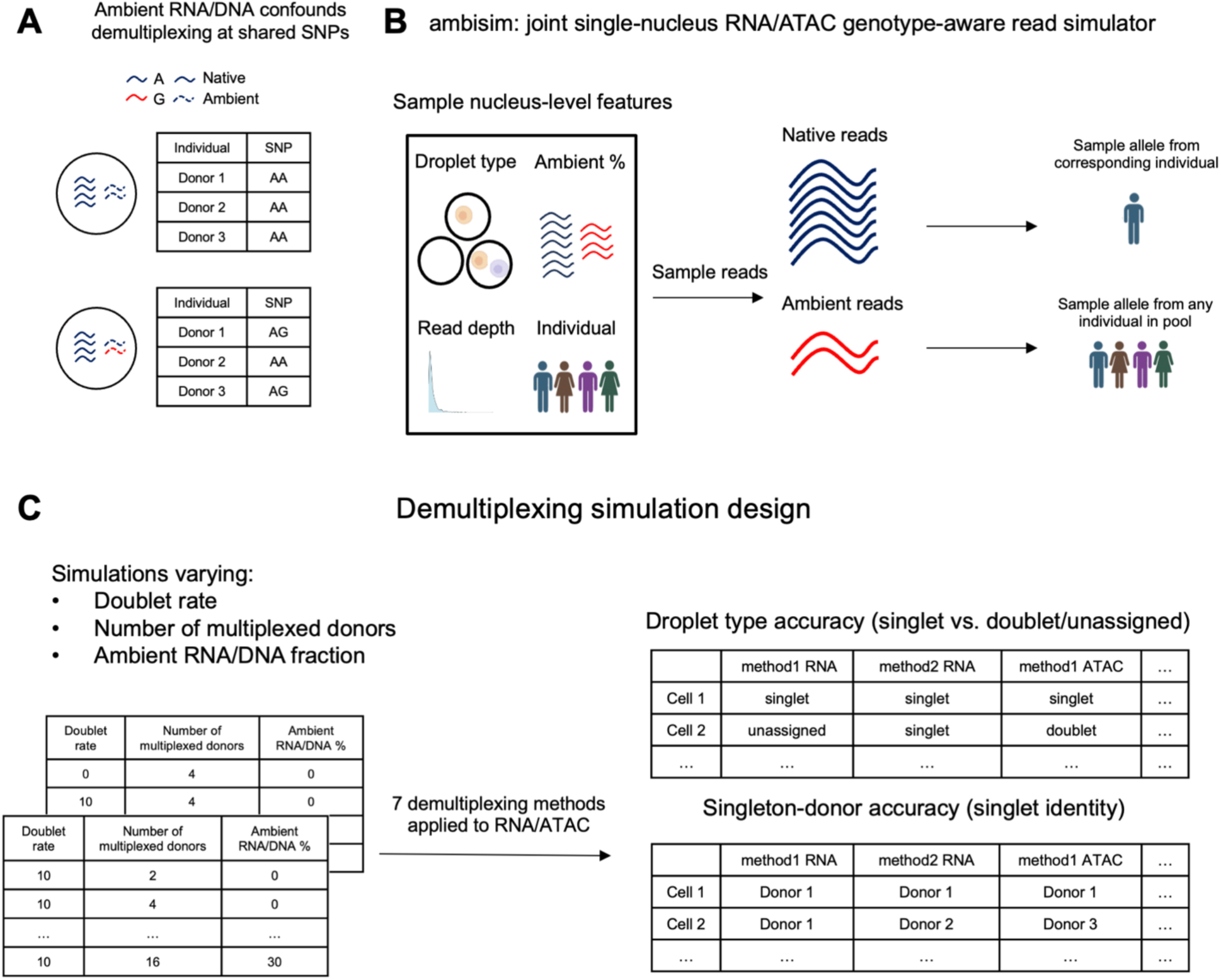
*ambisim*, a multiplexed joint snRNA/snATAC read simulator that controls ambient proportions. A) Example of a scenario in which ambient RNA can confound demultiplexing methods. B) Design of *ambisim.* Cell attributes, including donor identity and ambient RNA fraction, are specified, and then reads are sampled from a reference genome. For any read that overlaps a SNP, an allele is sampled from the donor’s genotype. If the read is an ambient read, a SNP is instead sampled based on all donors’ genotypes. C) Simulation designs. Heterotypic doublet proportion and number of multiplexed donors are varied along with ambient RNA/DNA fraction. Demultiplexing methods are evaluated based on two metrics: droplet type accuracy (percentage of droplets assigned to the correct droplet type) and singleton-donor accuracy (percentage of singlets assigned to the correct donor).

To model ambient RNA, previous studies adopted a partial simulation framework^10,13,15,22^. First, ground truth singlets are generated from *in silico* mixtures from independent single-donor experiments. Doublets are then artificially generated by merging reads across random pairs of singlets, and ambient RNA is induced by randomly swapping reads across barcodes according to a global contamination rate. However, this approach is problematic, as ambient material tends to originate from non-informative parts of the genome, such as regions outside of genes and peaks^26,27^. While single cell read simulators for both RNA and ATAC exist and can be used to control different experimental designs^28,29,30^, they are currently unable to model ambient material and cannot simulate from genotypes.

To address these issues, we developed *ambisim* to generate more realistic multiplexed single nucleus multiome data. In addition, we benchmarked seven genotype-based demultiplexing methods, three of which can be applied with or without reference genotypes **(Table S1, Supplemental Information)**, in a comprehensive set of *ambisim* simulated multiplexed snRNA/snATAC and two real multiplexed snRNA/snATAC datasets. We observed variable performance of methods as a function of study design and ambient contamination. Finally, we propose a new metric, *variant consistency*, which leverages cell-level allele counts to estimate ambient contamination, which we used to further characterize differences in demultiplexing quality between methods.

## Results

### *ambisim*: an ambient RNA/DNA aware joint RNA/ATAC read simulator

*ambisim* requires an input a reference joint RNA/ATAC dataset and a reference genotype VCF file. It then generates a droplet set and a corresponding set of fastq files encapsulating the data in a simulated multiplexed experiment. First, for each input barcode, *ambisim* samples a droplet type – singlet, doublet, or empty – and then samples cell-level attributes, including read depth, droplet type, individual identity, and an ambient read fraction **(Fig 1B)**. Then, *ambisim* samples reads from the genome according to probability distributions based on a read’s ambient status. If the read overlaps a SNP, an allele is sampled based on two cases. If the read is native to one individual, then an allele is sampled from that individual’s genotype. If the read is ambient, then the allele is sampled from all individuals’ genotypes according to the ambient distribution. Complete details on *ambisim* are provided in the Methods section.

We generated simulations across two sets of experiments. In the first set, we fixed the number of multiplexed donors to 4 and varied the doublet rate between 0% and 30%. In the second set, we varied the number of multiplexed donors between 2 and 16 and fixed the doublet rate to 10%. In total, we generated 44 simulated datasets **(Fig 1C, Table S2)**. In each experiment, we sampled reads across 9,000 droplets for a mean coverage of 25,000 reads for RNA and 40,000 reads for ATAC (according to the 10X Genomics sequencing depth recommendation) and multiplexed an equal proportion of each individual **(Table S3)**. After generating the simulated experimental data, we applied each demultiplexing method to both the RNA and ATAC BAM files independently, filtering for SNPs with imputation R^2^ > 0.90 for methods requiring reference genotypes **(Methods)**.

### Simulations reveal variable demultiplexing method performance as a function of ambient RNA/DNA

We evaluated demultiplexing methods in both modalities based on two metrics: droplet-type accuracy, and singleton-donor accuracy **(Fig 1C).** Droplet-type accuracy denotes the proportion of droplets that are identified correctly (singlet vs. unassigned), and singleton-donor accuracy denotes the proportion of singlets that are called the correct individual. We compared each method’s performance as a function of increasing levels of ambient RNA/DNA across both sets of simulations **(Figs 2A–2H, Figs S1–2).** As expected, we observed that ambient contamination impacted the performance of most methods. When considering droplet-type accuracy, we found that the increase in contamination generally led to stable decreases in accuracy. However, when considering singleton-donor accuracy, we found that several methods, primarily genotype-free ones, were unstable across ambient contamination estimates across both modalities. We note that freemuxlet was not designed for high-coverage data and would likely perform similarly to other genotype-free methods with minor modifications.

**Figure 2:**
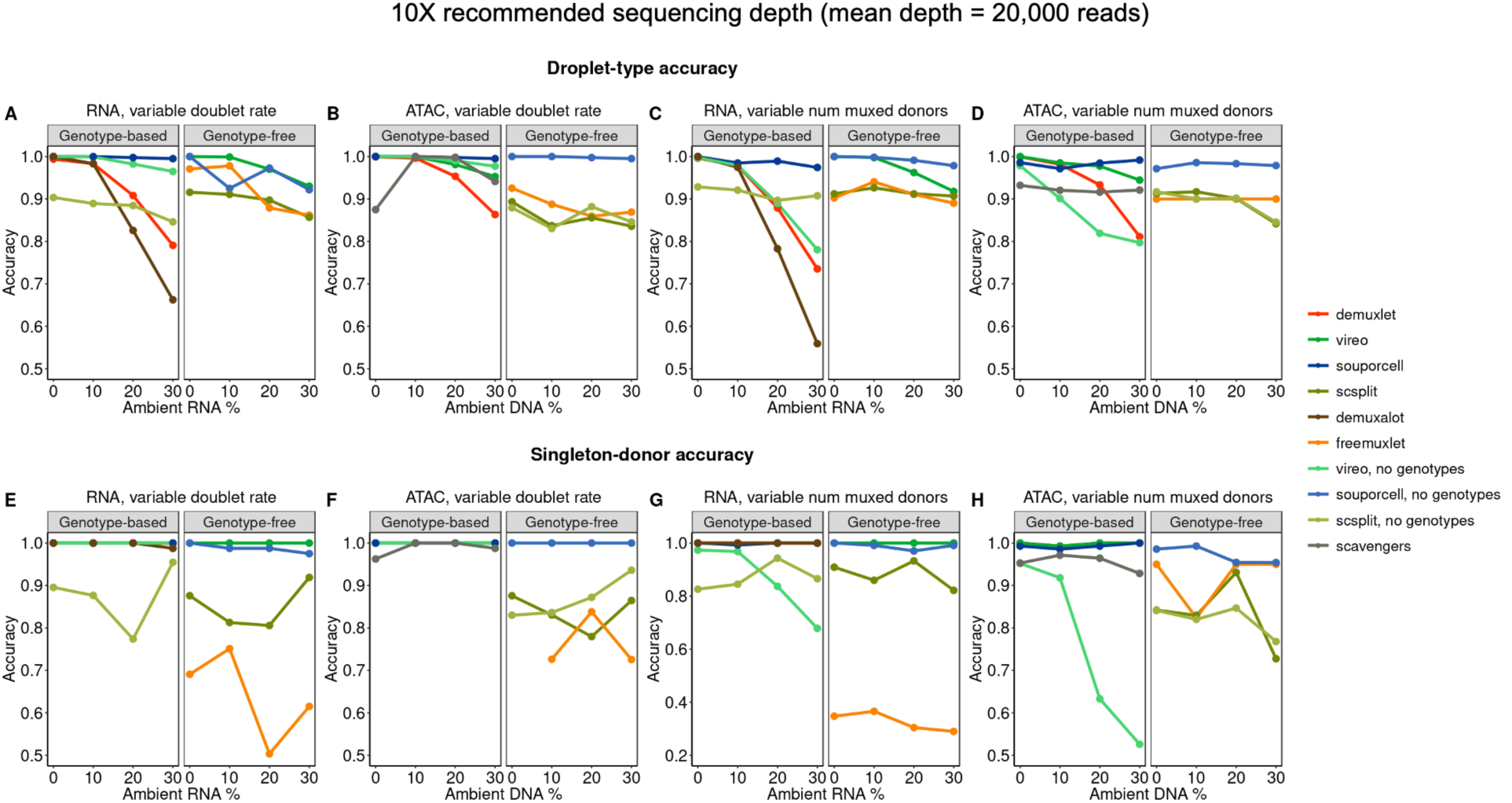
Accuracy comparisons in simulations. A) Comparison of droplet-type accuracy as a function of ambient RNA/DNA, summarized across experiments with number of multiplexed donors = 4 and variable doublet rates. B) Same as A), but for ATAC. C) Comparison of singleton-donor accuracy as a function of ambient RNA/DNA, summarized across experiments with doublet rate = 10% and variable numbers of multiplexed donors. D) Same as C, but for ATAC. E) Same as A), but for singleton-donor accuracy. F) Same as B), but for singleton-donor accuracy. G) Same as C), but for singleton-donor accuracy. H) Same as D), but for singleton-donor accuracy.

We next asked how methods performed on average across various parameter settings. We found that both doublet rate and the number of multiplexed donors had modest impacts on many methods, with some methods’ performance being disproportionately affected (eg. freemuxlet for doublet rate, vireo no genotypes for number of multiplexed donors) **(Fig S3).** When evaluating demultiplexing methods between modalities, we found that many ATAC-based methods performed slightly better than their RNA-based counterparts, **(Fig S4).** Finally, we found that genotype-based methods perform modestly better than genotype-free methods **(Fig S5).**

We next asked if there were common properties among droplets misidentified by each method. When examining whether inaccurate droplets tended to be miscalled as singlets or doublets, we found differences between methods – for example, genotype-based methods, such as demuxlet and demuxalot, identify more doublets, while most genotype-free methods identify more singlets **(Fig S6)**. Between accurate and inaccurate droplets, we found that the distribution of the number of SNPs was different, for most methods **(Fig S7)**. When considering the true ambient RNA/DNA distributions, we found a similar trend among most genotype-based methods: misclassified droplets tended to have higher ambient contamination **(Fig S8)**. However, for several genotype-free methods, such as freemuxlet and scSplit, the distributions of ambient RNA/DNA between accurate and inaccurate droplets were smaller, suggesting that latent genotype inference is the driving force.

### Many demultiplexing methods perform similarly in lower sequencing-depth simulations

One possible design choice in droplet-based single-nucleus experiments is to sequence more nuclei, at the cost of a lower per-nucleus sequencing depth. This may be an advantageous design choice for several reasons, such as increasing the likelihood of sampling rare cell types and increasing the statistical power of quantitative trait loci mapping by identifying more nuclei per individual. To measure the effects of sequencing depth on demultiplexing method accuracy, we generated an identical set of simulations with a lower depth (mean 7,000 reads for both assays) (**Fig 3A–3G, Figs S9–10, Table S4).** We found that, similar to the higher-coverage simulations, demultiplexing method performance decreases stably across ambient contamination levels, with higher variance for methods such as demuxlet and demuxalot. In general, many methods performed similarly across coverage levels, but singleton-donor accuracy in ATAC-based genotype-free methods is disproportionately affected by coverage **(Fig S11).** One likely explanation for this behavior is that the number of covered SNPs scales with sequencing depth differently for RNA and ATAC, as ATAC reads are derived from a larger portion of the genome, which results in SNPs being more sparsely covered **(Fig S12)**.

**Figure 3:**
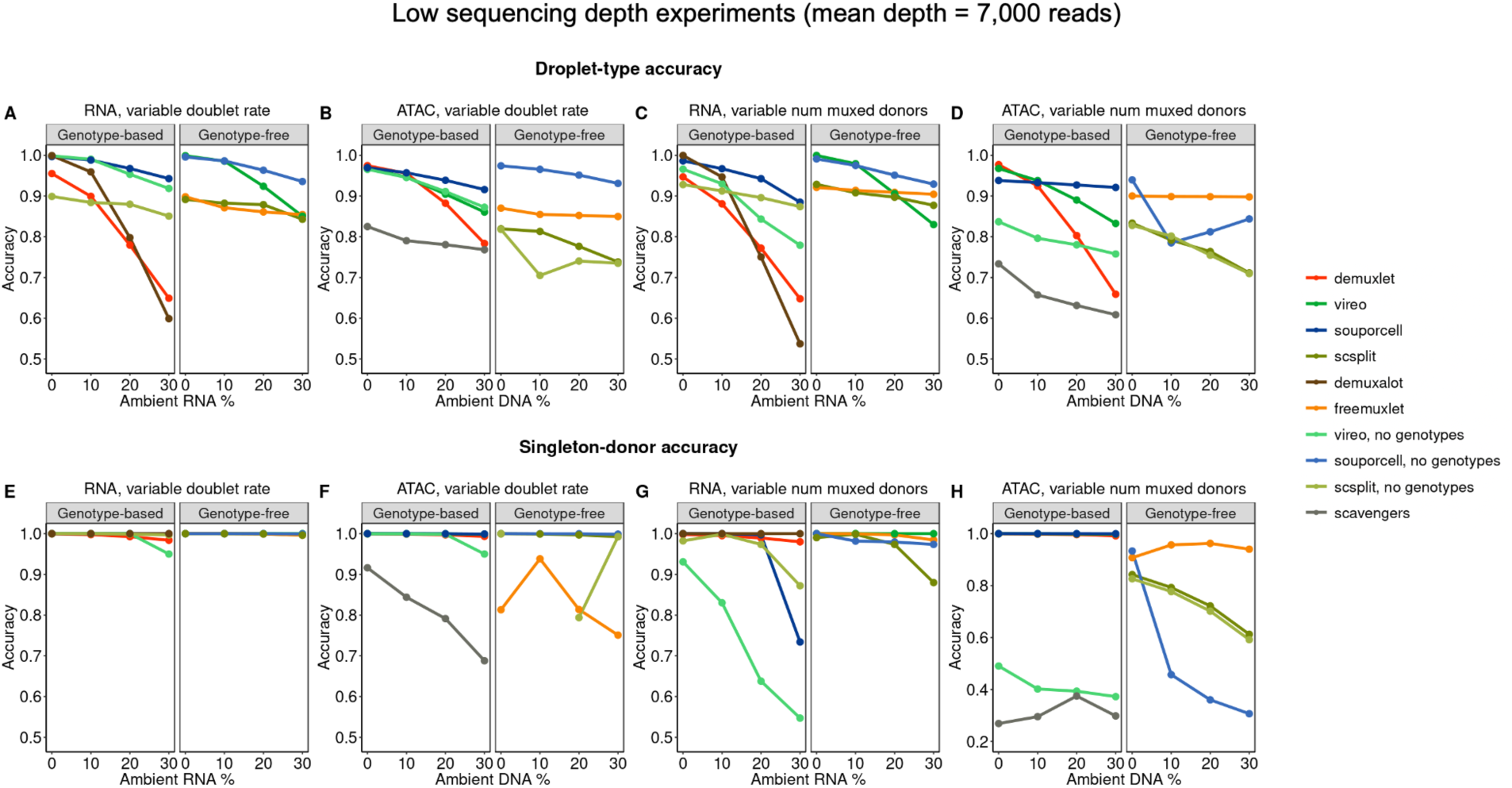
Accuracy comparisons for lower-coverage versions of simulations. A) Comparison of droplet-type accuracy as a function of ambient RNA/DNA, summarized across experiments with number of multiplexed donors = 4 and variable doublet rates. B) Same as A), but for ATAC. C) Comparison of singleton-donor accuracy as a function of ambient RNA/DNA, summarized across experiments with doublet rate = 10% and variable numbers of multiplexed donors. D) Same as C, but for ATAC. E) Same as A), but for singleton-donor accuracy. F) Same as B), but for singleton-donor accuracy. G) Same as C), but for singleton-donor accuracy. H) Same as D), but for singleton-donor accuracy.

### Application of demultiplexing methods to real joint snRNA/snATAC data reveals low between-method droplet-type correlations

We next applied demultiplexing methods to two multiplexed joint snRNA/snATAC datasets to evaluate their consistency across within and across experiments. The first was a 10X Multiome dataset collected across seven runs in aorta samples (n=35,201 droplets, multiplexed across 12 individuals)^20^. The second was an in-house multiplexed induced pluripotent stem cell (iPSC) reprogramming experiment (n=30,497 droplets, multiplexed across 4 individuals), with comparable coverage levels between datasets **(Fig S13).** When comparing the donor assignment distributions between methods, we found that many methods identify similar proportions of singlets and non-singlets, with the exception of freemuxlet consistently identifying more singlets in both modalities **(Fig 4A–B, Tables S5–S6).**

**Figure 4:**
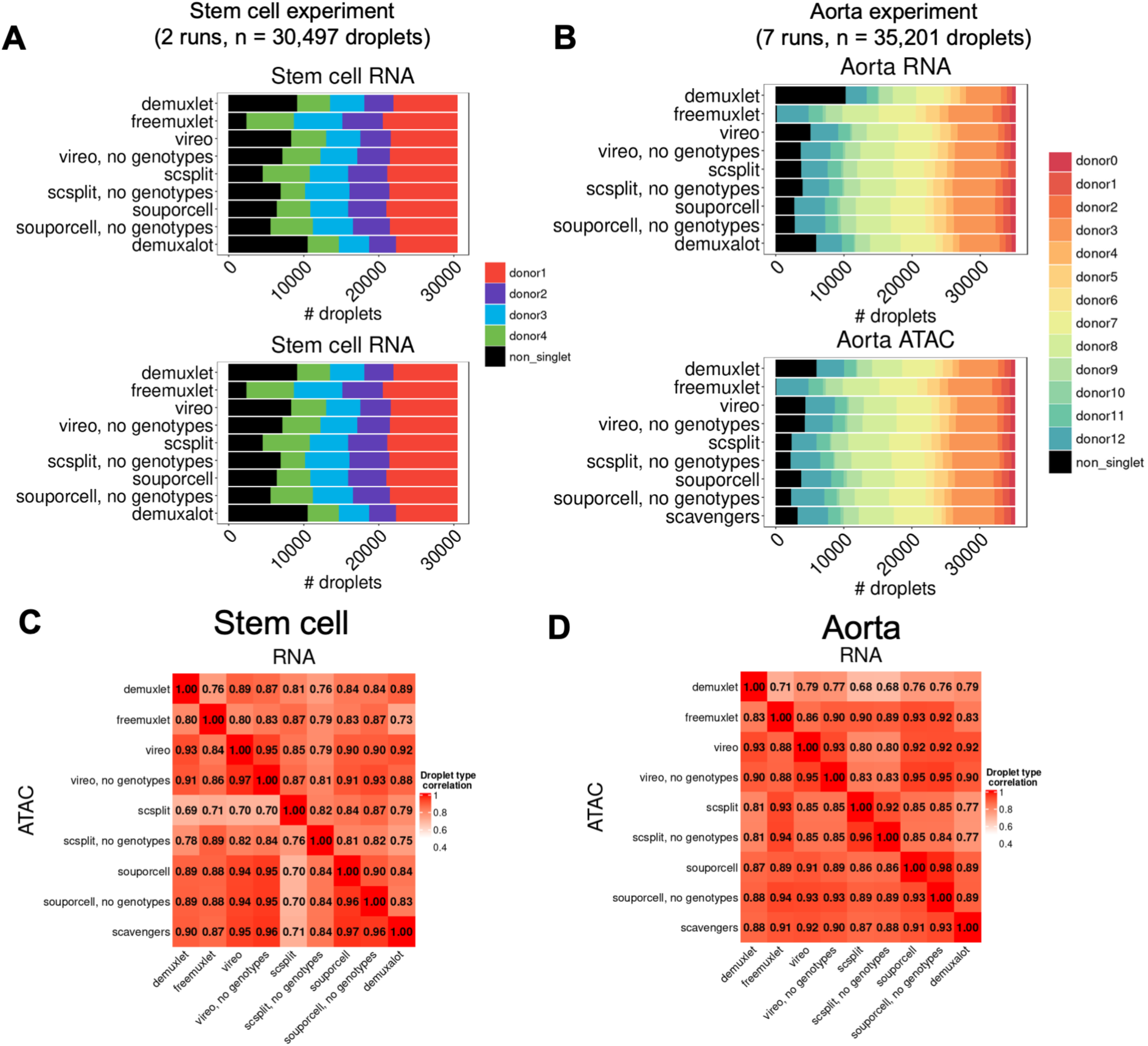
Comparing demultiplexing within modalities in real data. A) Distributions of singlet and doublet calls in stem cell dataset. B) Same as A), but for aorta dataset. C) Droplet-type correlations between methods in RNA (top) and ATAC (bottom) in stem cell dataset. D) Same as C), but for aorta dataset.

We next asked how the demultiplexing output differed between methods by comparing correlations of droplet type and singleton-donor assignment. Within each modality, droplet-type correlation was high for most pairs of methods **(Fig 4C–D)**, but the range was variable (0.69– 0.97 stem cell, 0.71–0.93 aorta). Singleton-donor correlation was also high for most pairs of methods (0.99–1.00 stem cell, 0.96–1.00 aorta), except for scSplit in both RNA and ATAC (0.89–1.00 stem cell, 0.85–0.90 aorta) **(Fig S14)**. We noticed that our between-method droplet-type correlations were lower than previously reported in scRNA-seq^21^, which may be due to increased contamination in single-nucleus data. When comparing droplet-type correlations between a representative simulation from *ambisim* (num donors = 4, doublet rate = 20%, ambient rate = 20%) and the iPSC reprogramming experiment, we found that while between-method correlation patterns were similar, the correlation in the iPSC experiment was lower compared to the simulation **(Fig S15).**

Recently, there has been interest in using ensemble methods combining multiple genotype-based demultiplexing methods^23,24^. Given our observation of lower between-method correlations, we next asked if it would be feasible to apply a similar ensemble framework to joint snRNA/snATAC data. To address this, we examined how concordant the singlet sets were across modalities and multiple methods. The cross-modality intersection indicated that the average between-method pairwise intersection is similar within and across modalities **(Fig 5A)**. However, the droplet-type intersection between multiple methods was much lower (0.52–0.73 stem cell, 0.46–0.71 aorta) **(Fig 5B)**, while the individual intersection remained high (0.94–0.99 stem cell, 0.72–0.83 aorta). An UpSet visualization of droplet-type intersections between methods revealed that droplets outside the all-method intersection do not arise from specific methods, but rather combinations of methods **(Fig 5C–D).** These results suggest that single-nucleus data presents additional challenges for demultiplexing compared to single-cell data, and that the intersection of multiple methods may not yield coherent demultiplexing sets. In the next section, we propose and apply a new metric to compare demultiplexing methods by estimated ambient contamination.

**Figure 5:**
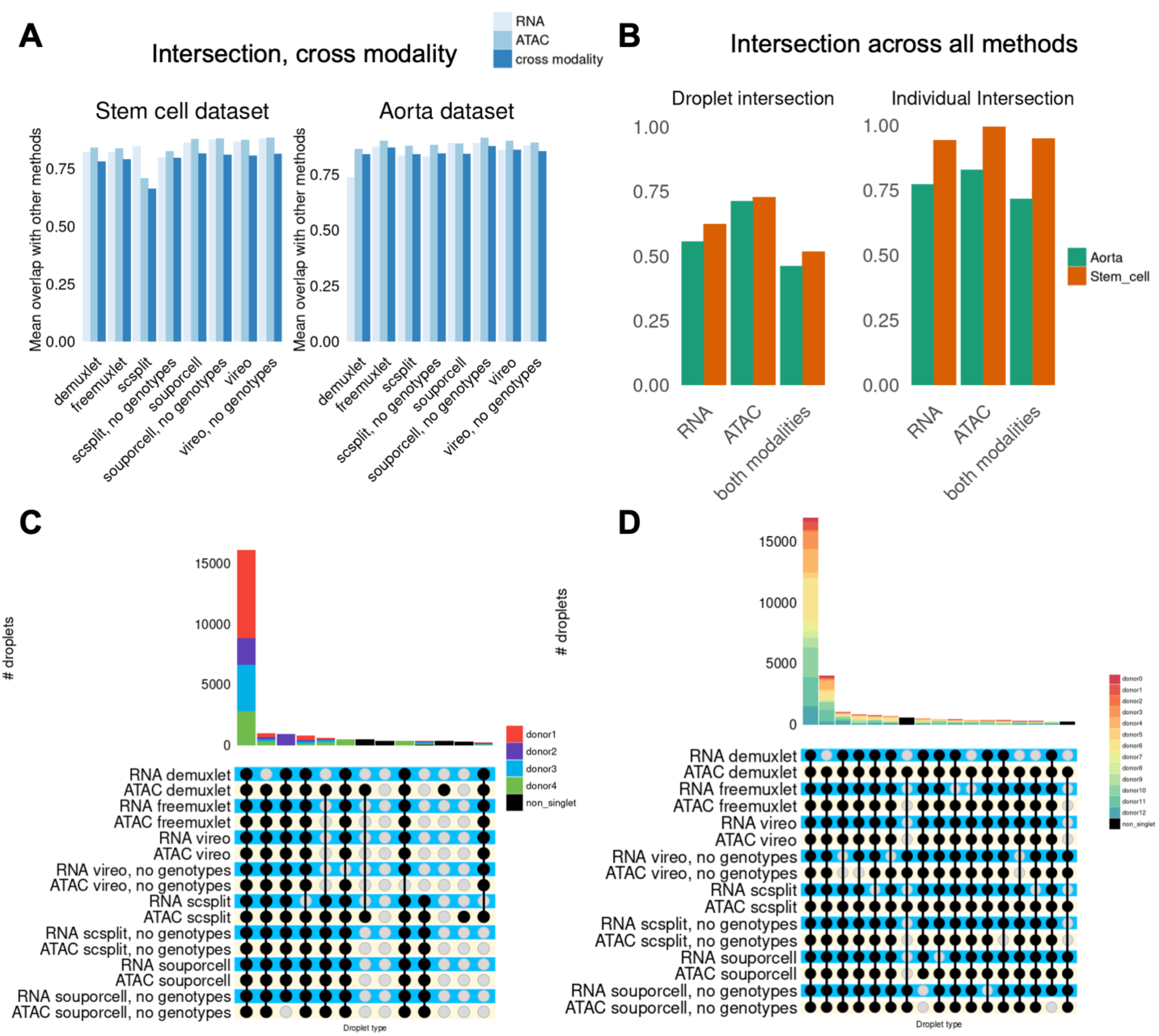
Comparison of demultiplexing methods within and across modalities between multiple methods. A) Mean droplet-type overlap between methods, including across modalities. B) Proportion of nuclei that are called the same droplet type and individual across all methods, both within and across modalities. C) UpSet visualization of all methods with both RNA/ATAC-based demultiplexing in the stem cell dataset. D) Same as C), but for the aorta dataset.

### Variant consistency, a metric to compare demultiplexing methods by estimated ambient contamination

We reasoned that reference genotypes could be leveraged post-hoc to provide an alternative evaluation of demultiplexing method quality. By quantifying how often a nucleus’s variant set agrees with their assigned donor’s genotype, we can estimate residual ambient RNA/DNA rates on a per-method basis. We define this metric as *variant consistency*. Our variant consistency framework requires the following as input: per-nucleus individual assignments, a pileup matrix of allele counts, and a genotype VCF file. We then classify allele counts of variants as consistent or inconsistent based on whether they match the assigned individual’s genotype **(Fig 6A, Methods)**. We divide consistency counts into sub-categories. Variants that are consistent only in one individual are denoted C1 variants. Variants that are consistent in multiple individuals are denoted as C2 variants. Variants that are inconsistent in one individual, but consistent in at least one other individual are denoted as I1 variants. Finally, variants that inconsistent across all individuals are denoted as I2 variants. Each of these categories provides different information to characterize variant-level properties. I1 reads represent the lower bound of ambient contamination, as those reads must originate from another donor. I2 variants represent sequencing or genotyping errors, as those allele counts do not align with any individual’s genotypes. Our subsequent analyses focus on the difference in the number of I1 read counts between methods and modalities.

**Figure 6:**
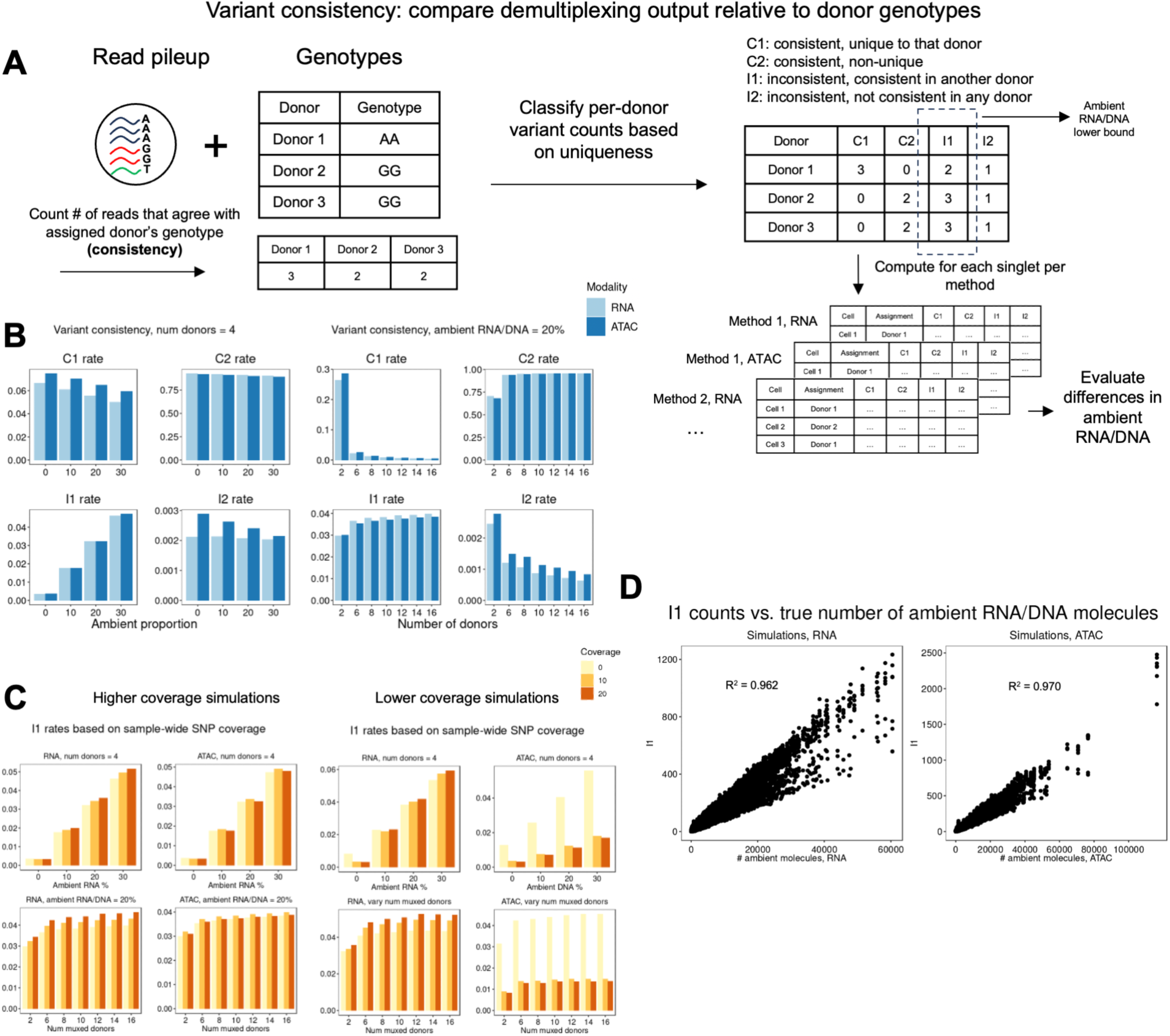
Concept of variant consistency metric and application to simulations. A) Schematic of *variant consistency* metric. Per nucleus allele counts are classified into four categories of consistency based on the uniqueness of each variant. B) Variant consistency ratios based on ambient RNA rates and multiplexed donors in simulated experiments. C) I1 rates based on how many times a SNP is covered experiment wide in lower-depth simulations. D) Correlation of allele counts that are inconsistent but non-unique (I1) with true ambient RNA/DNA across simulations is high, showcasing our ability to quantify ambient contamination in real datasets.

We first applied our metric to the simulated experiments and compared the distributions of variants across ambient proportions **(Table S7)**. When comparing consistency counts for the same nuclei across both modalities, we noticed that the C1 rates are higher in ATAC than RNA, which suggests that more informative variants lie in non-coding regions of the genome **(Fig 6B)**. Additionally, C1 rates rapidly decrease based on the number of multiplexed donors. We found that in our lower-coverage simulations, I1 rates of SNPs vary based on how many times that SNP is covered within an experiment **(Fig 6C).** When comparing cell-level I1 counts to their corresponding true number of ambient RNA/DNA molecules, we found that the correlation is higher in ATAC (R^2^ = 0.970) compared to RNA (R^2^ = 0.962) **(Fig 6D)**.

### Variant consistency shows variable demultiplexing method uncertainties in assignments

We next applied our variant consistency framework on real data to compare demultiplexing method performances. We first subsetted singlets to the ones that are uniquely identified by each method, and found high variance, particularly in freemuxlet and scSplit no genotypes across RNA/ATAC **(Fig 7A)**. We ran our framework on the unique droplets called by each method and compared C2 and I1 rates **(Fig 7B, Tables S8–S9)**. Notably, I1 rates differed across methods in both modalities – scSplit called the most contaminated singlets, while demuxalot and demuxlet called the least contaminated singlets. Across datasets, the contamination patterns between RNA and ATAC were inverted, highlighting the utility of our variant consistency metric in understanding ambient contamination patterns on a per-dataset basis.

**Figure 7:**
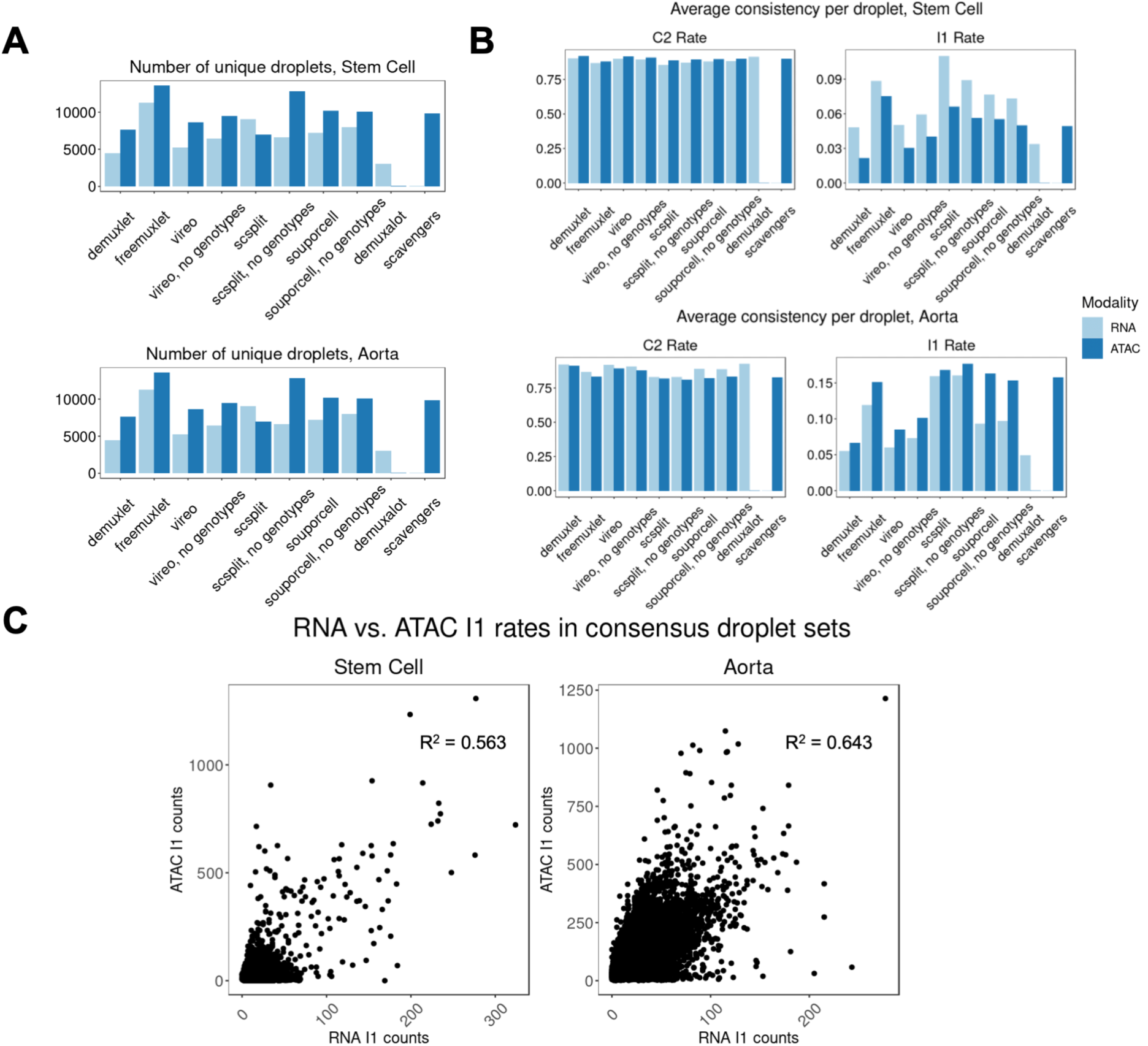
Applying variant consistency to real data reveals differences in demultiplexed singlets quality between methods in both RNA and ATAC. A) Counting the number of droplets that each method detects outside of the within-modality intersection. B) C2 and I1 rates in the droplets called uniquely by each method. C) Correlation of I1 rates between modalities.

When comparing contamination rates alone, we found that demuxlet, vireo, and demuxalot called droplets with the lowest amount of contamination. However, when considering the number of unique droplets identified by each method, demuxlet (RNA n=4,453, ATAC n=7,598) and demuxalot (RNA n=3,029) were the most conservative. Finally, when comparing consistency distributions per donor in both experiments, we found that variant consistency rates are similar across individuals, except for freemuxlet RNA **(Figs S16–S17)**, suggesting that demultiplexing methods in these datasets do not have biases towards particular individuals.

## Discussion

In this study, we introduced ambisim, a joint snRNA/snATAC read-level simulator that can control ambient molecule proportions and sample from reference genotypes, and a novel metric, *variant consistency*, which enables direct comparison of demultiplexing methods based on the quality of the identified singlets. Using these tools, we compared the performance of seven demultiplexing methods in simulated and real multiome data. Our simulations showed that varying levels of ambient RNA/DNA present problems for demultiplexing methods, which would therefore benefit from directly modeling ambient material. In real data, we found that variable between-method correlations suggesting that ensemble methods may not be reliable in single-nucleus experiments in both RNA and ATAC. Finally, our variant consistency metric provides a framework for estimating ambient RNA/DNA proportions in real data and comparing demultiplexing uncertainty between methods.

## Methods

### Aorta endothelial data

Fastq files were mapped using Cellranger-Arc v.2.0.1 with default parameter settings. VCF files were imputed using the TopMed imputation server using R^2^ > 0.3 and the 1000 Genomes Phase 3 Reference Panel^34,35,36^.

### Stem cell data processing

10X Multiome data was mapped to 10X Genomics’ pre-set hg38 reference using CellRanger Arc v.2.0.1 with default parameter settings. Cellranger BAM files and peaks were subset to autosomal chromosomes.

### Demultiplexing algorithms

#### Demuxlet/Freemuxlet

Demuxlet is a maximum-likelihood method that requires genotypes as input to the method. After counting alleles at each SNP in a reference set, it uses a statistical model which incorporates base error probabilities and models allele probabilities using a binomial distribution. It also extends the statistical model designed for singlets to detect doublets by modeling all combinations of possible doublets multiplexed in the experiment. The ‘dsc-pileup’ command in the popscle software suite was used to generate read pileups for each BAM. Then, ‘popscle demuxlet’ was run with default options.

#### cellSNP/Vireo

Vireo uses a variational inference approach to estimate latent genotypes. It directly includes the option of including genotypes to establish priors on the genotypes. cellSNP-lite^31^ was used to generate allele counts per cell based on a variant reference set. Per the developers’ recommendations, cellSNP-lite was applied to a common variant set derived from 1000 Genomes data^44^ filtered with a minor allele frequency > 0.01, and SNPs with coverage < 20 across the run were discarded. Vireo was then run with and without genotypes based on the cellSNP-lite output.

#### scSplit

scSplit uses an expectation-maximization approach and identifies distinguishing variants to minimize the number of variants used for demultiplexing. The pipeline consists of filtering reads, calling variants directly from the data using freebayes^32^, allele count pileup, and model fitting. Genotypes were supplied as priors for the model during the final demultiplexing step. A modified version of scSplit was used to reduce computational time. Briefly, the cells by variants matrix was computed using cellSNP-lite, and a custom version of scSplit designed for reading in cellSNP-lite input was used.

#### Souporcell

Souporcell is a sparse mixture model that clusters cells according to allele fractions. The cluster assignments are optimized using a simulated annealing algorithm. For the genotype-free version, souporcell was supplied with the same 1000 Genomes variant set as previously mentioned for the --common_variants parameter. Souporcell no genotypes was tested without including the common-variants parameter, in which variants are called directly from the data using freebayes, but the performance was determined to be consistently better when including common variants.

#### Demuxalot

Demuxalot is a Bayesian method that aggregates SNPs across genes. It calls variants, assigns base frequencies, and alternately estimates genotype signatures and assignments. Demuxalot was unable to be applied to ATAC-seq due to the lack of UMIs.

#### scAVENGERS

scAVENGERS is a method developed for multiplexed scATAC-seq experiments. It derives reference and alternate allele probabilities by modeling pre-PCR amplification counts based on ATAC barcode attachment probabilities. Strelka2^33^ was used to call variants from the data, and Vartrix was used to generate the cells by allele counts matrix. A modified version of the scAVENGERS pipeline was then used to derive cell assignments.

### Variant consistency

Given a multiplexed single-cell experiment with *D* droplets, *V* covered variants, and *K* individuals, we are interested in computing both the consistency of each variant between the read pileup and the assigned donor’s genotype and the total consistency proportions in each cell. More specifically, the input to this framework is matrices *R* ∊ *N^D^*^×*V*^, *A* ∊ *N^D^*^×*V*^, and genotype array *G* ∊ *N^K^*^×*V*^ which correspond to the sparse reference and alternate counts matrices and genotyping array for *k*individuals. For each droplet *d* ∊ *D*, we extract the reference and alternate allele counts from *P* ∊ *V* variants, and for the genotype of individual *k* at variant *v*, denoted as *G*(*v*), we compute consistency counts consistent(*v*) for each variant, defined as follows:

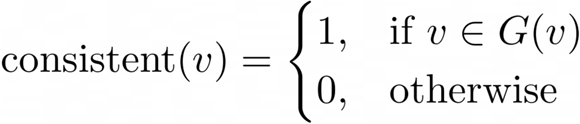

The output of the above framework is *k* consistency matrices, with each matrix *C* ∊ *N^D^*^×*V*^. Finally, we classify the reads into four categories: consistent with only that donor’s genotype (C1), consistent but matches another donor’s genotype (C2), inconsistent but matches another donor’s genotype (I1), and inconsistent and doesn’t match any donor’s genotype (I2). The final variant consistency matrix is *VC* ∊ *N^D^*^×*V*×*K*×4^. For each demultiplexing method, the output is a cell assignment matrix *Z* ∊ *N*^*D*×1^. Using *Z*, we iterate through *D* to extract the consistency counts corresponding from *VC* to the assigned individual, producing the final matrix *F* ∊ *N*^*D*×4^.

## Declarations

### Availability of data and materials

Raw fastq and VCF files for the aorta dataset were downloaded from GEO Accession GSE228428. Data for the in-house generated iPSC experiment will be available from the Impact of Genomic Variation on Function (IGVF) Consortium: data.igvf.org. Processed demultiplexing output is available on [Zenodo](https://zenodo.org/records/14827990?preview=1&token=eyJhbGciOiJIUzUxMiJ9.eyJpZCI6IjUzNzE5MzQ1LWFlMmItNDdiNi04NzhjLTY2NGQyN2M2ZDkzZCIsImRhdGEiOnt9LCJyYW5kb20iOiIwYTE5MTc3NTk1MGQzZTYzZjQ0OGEwZWZkOGFhOGRhYyJ9.-_KDCqFYmnZeoJVBsl5fMI2Db1fqbA9sRKaewY_Pjyqpzt1nTEtrZtuRDSZe5Q2XsUbP06VbKXpYkMJu7zY1sA).

Ambisim is available at https://github.com/marcalva/ambisim. The variant consistency pipeline is available at https://github.com/terencewtli/variant_consistency. All code used for generating data and figures, including mapping, demultiplexing, and simulation, is available at https://github.com/terencewtli/demux_benchmark_manuscript/.

### Authors’ contributions

T.L., M.A., and N.Z. conceived this project. T.L. performed the analysis and wrote the manuscript with guidance from N.Z and C.Y.L. M.A. designed and wrote the simulation software. C.Y.L. and N.Z. supervised the study. K.A., Y.S., and K.P. generated the stem cell reprogramming experiment. All authors read and edited the manuscript.

## Acknowledgments

We thank members of the Luo and Zaitlen labs for their feedback on the manuscript.

