## Supplemental Information for "The impact of ambient contamination on demultiplexing methods for single-nucleus multiome experiments"

#### Supplemental Figures

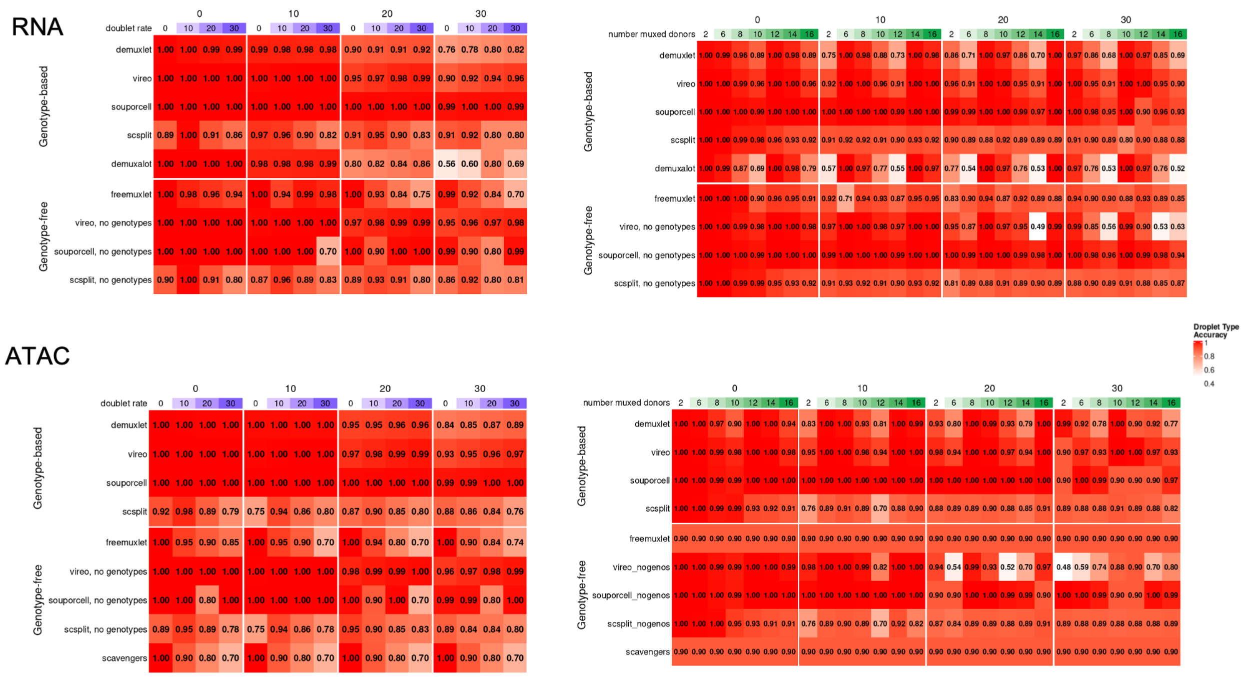


**Figure S1:** Full comparison of droplet-type accuracy across all simulations.

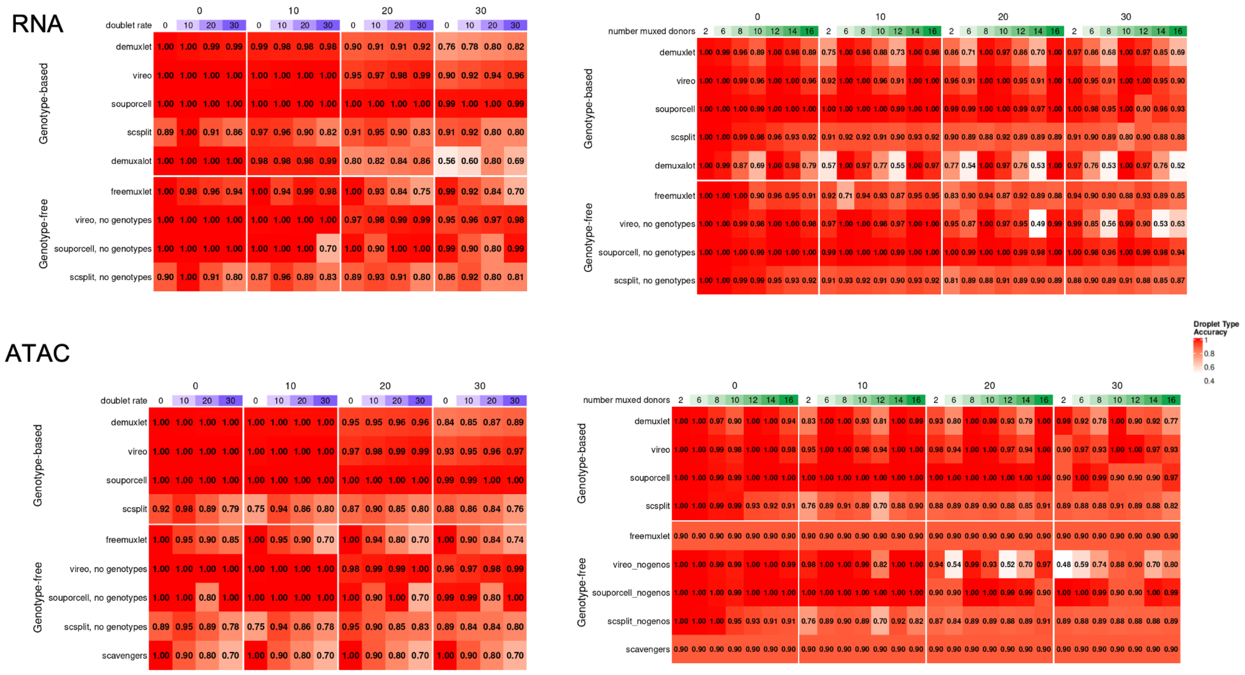


**Figure S2:** Full comparison of singleton-donor accuracy across all simulations.

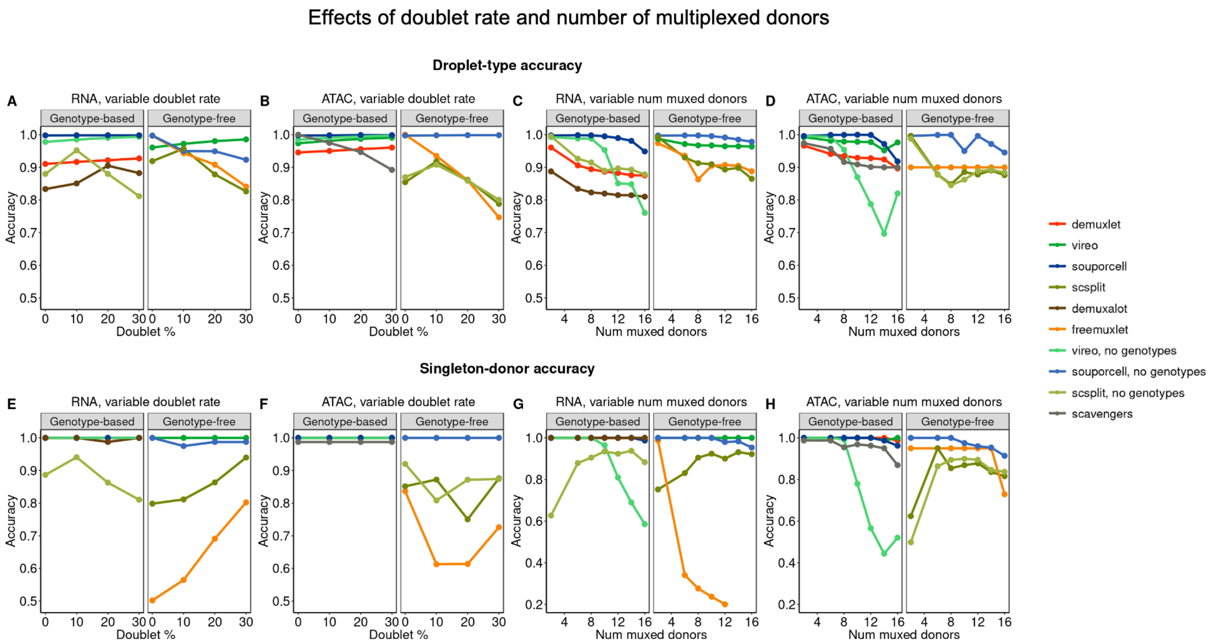


**Figure S3:** Effect of doublet rate and number of multiplexed donors on both accuracy metrics in full simulated experiments.

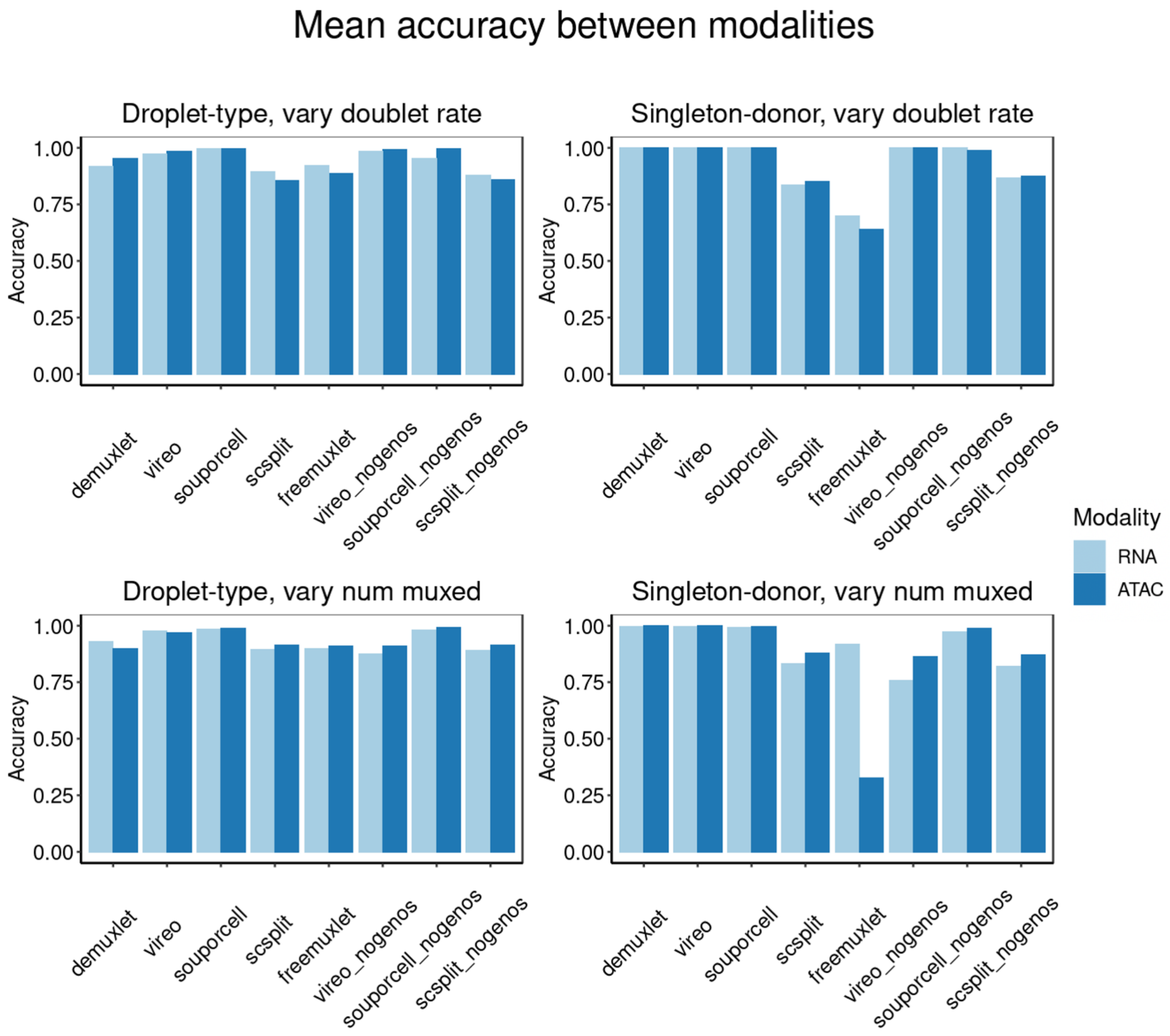


**Figure S4:** Mean accuracy across simulations between RNA and ATAC.

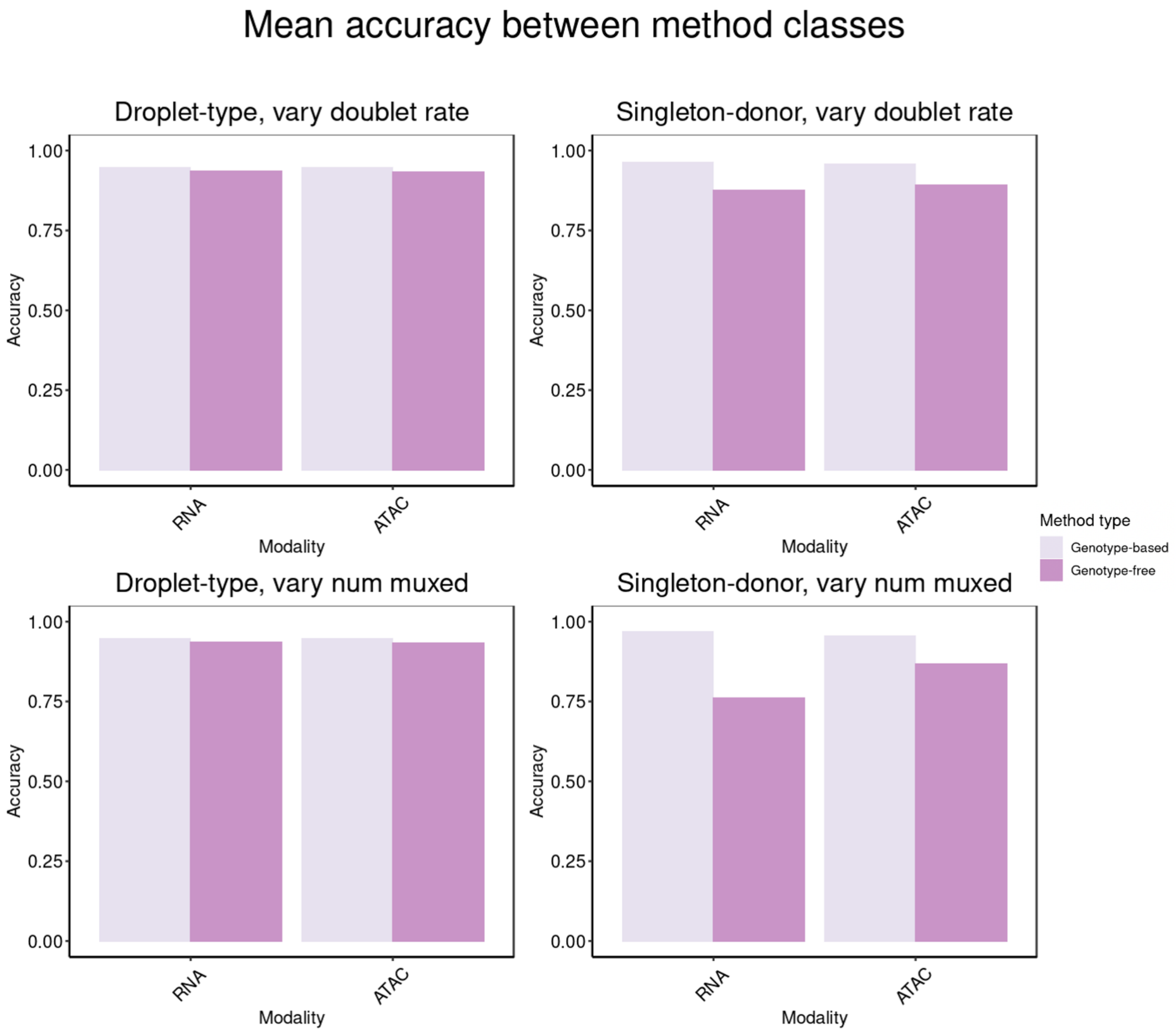


**Figure S5:** Mean accuracy between genotype-based and genotype-free methods across all simulations.

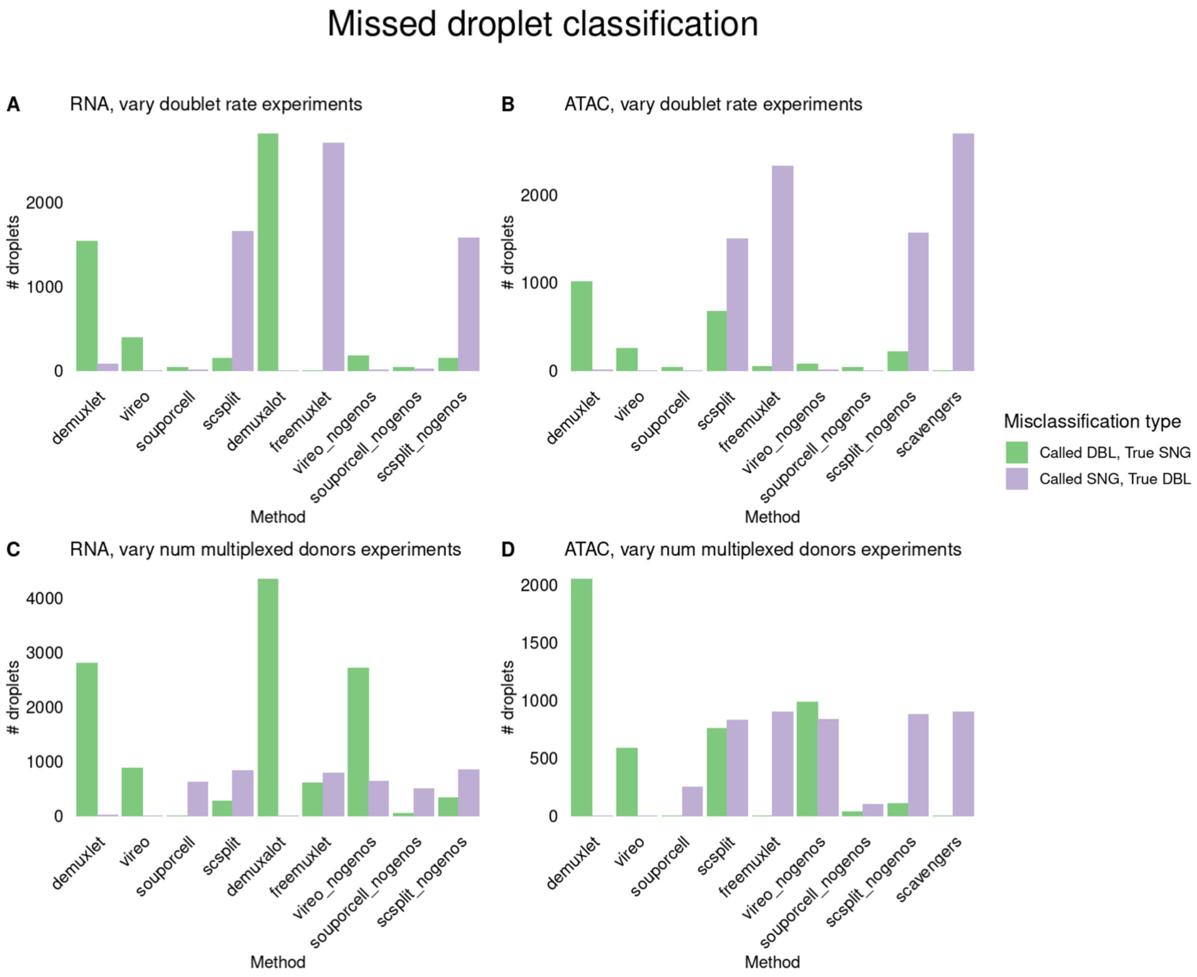


**Figure S6:** types of misclassified droplets across all simulations per method.

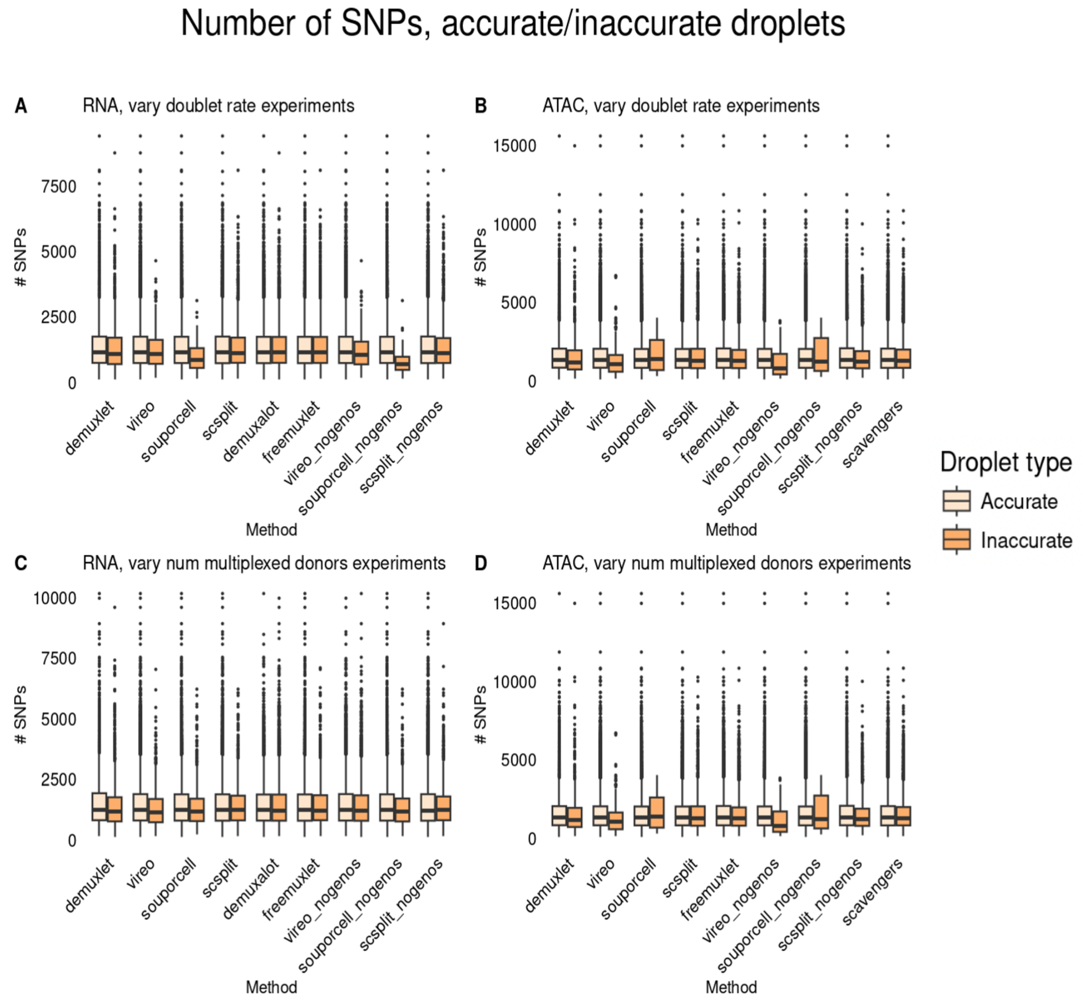


**Figure S7:** Comparing distributions of SNPs in uncontaminated experiments between accurate and inaccurate droplets.

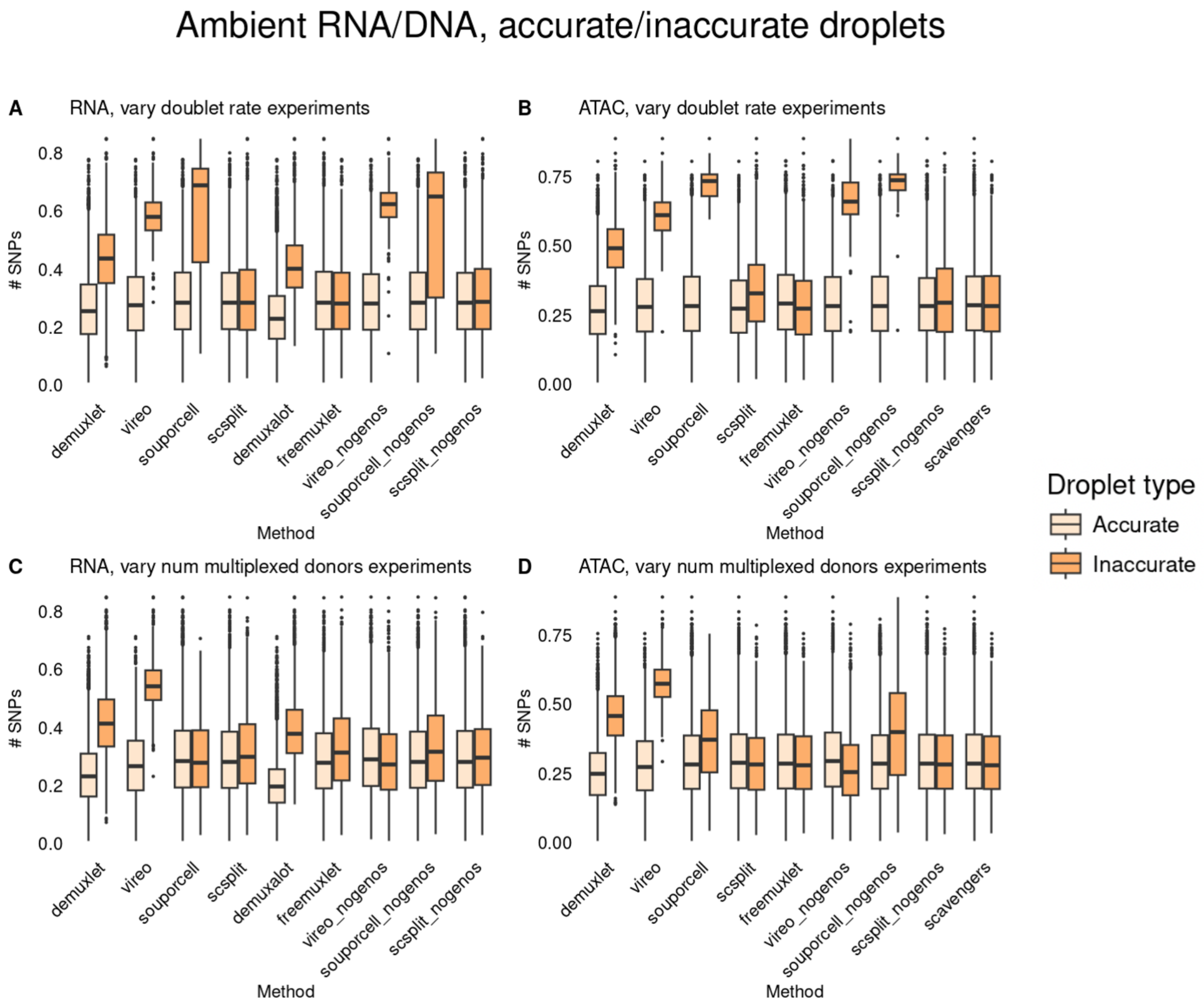


**Figure S8:** Comparing distributions of ambient RNA/DNA in cells called accurately and inaccurately in uncontaminated experiments.

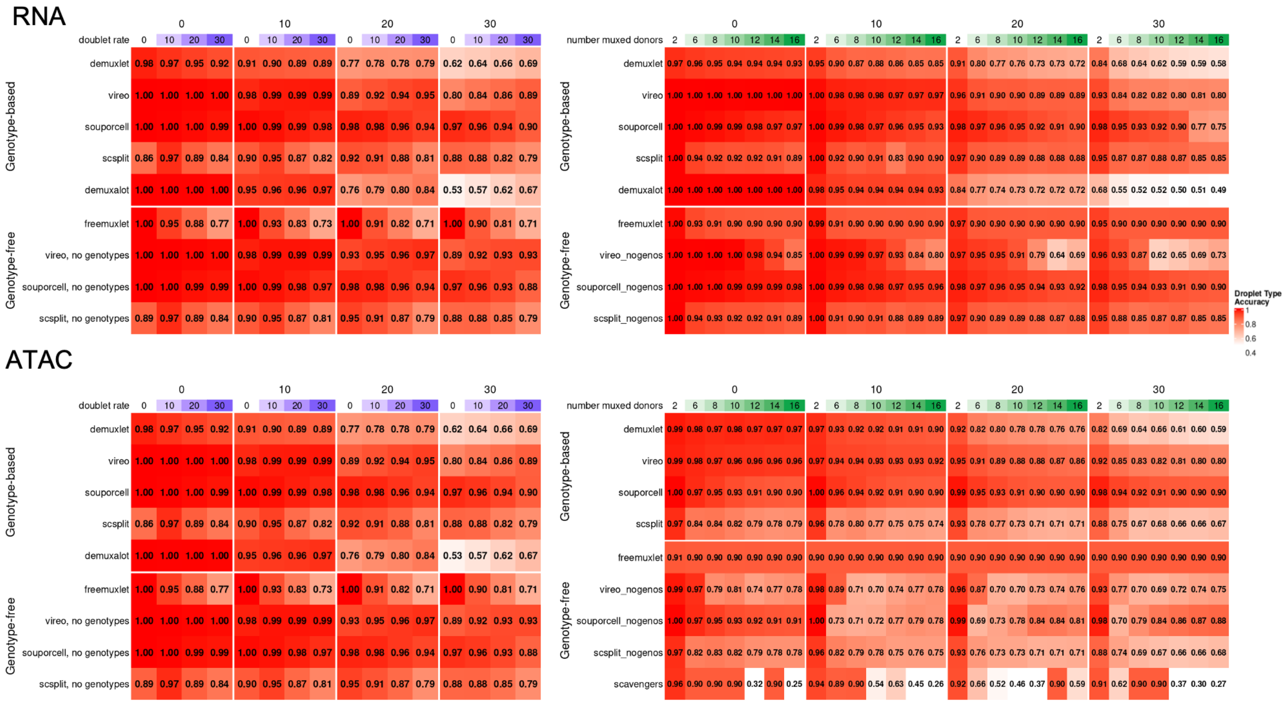

**Figure S9:** Droplet-type accuracy comparisons in lower-coverage versions of simulated experiments.

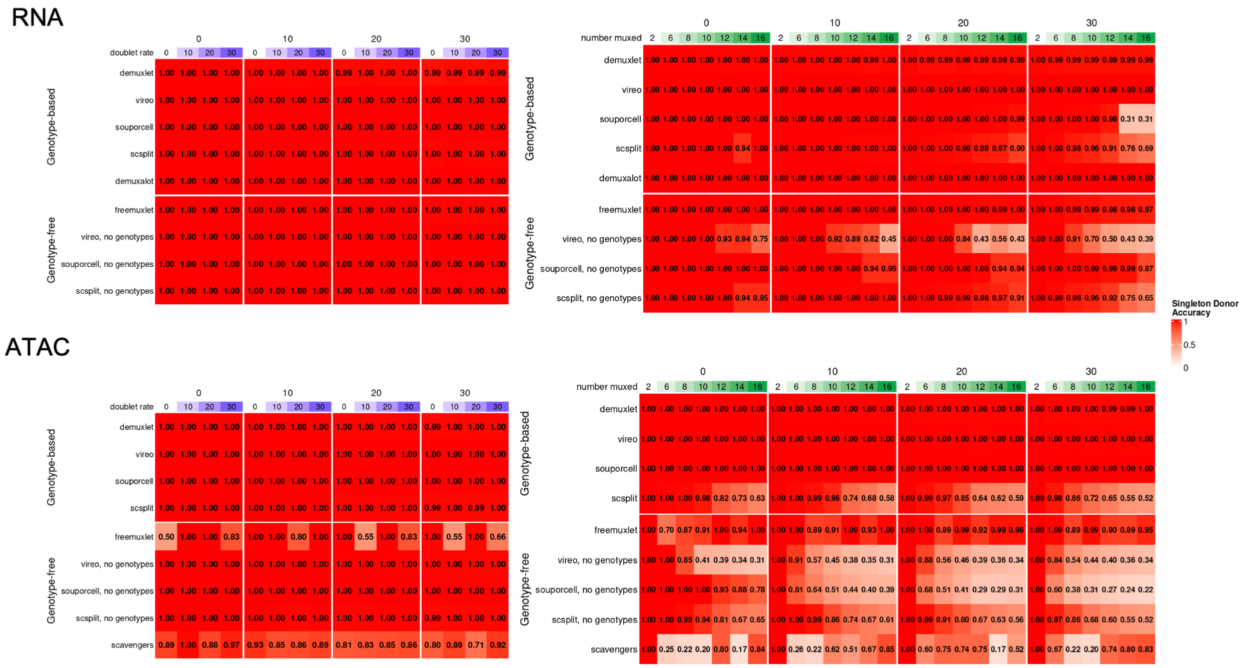


**Figure S10:** Singleton-donor accuracy comparisons in lower-coverage versions of simulated experiments.

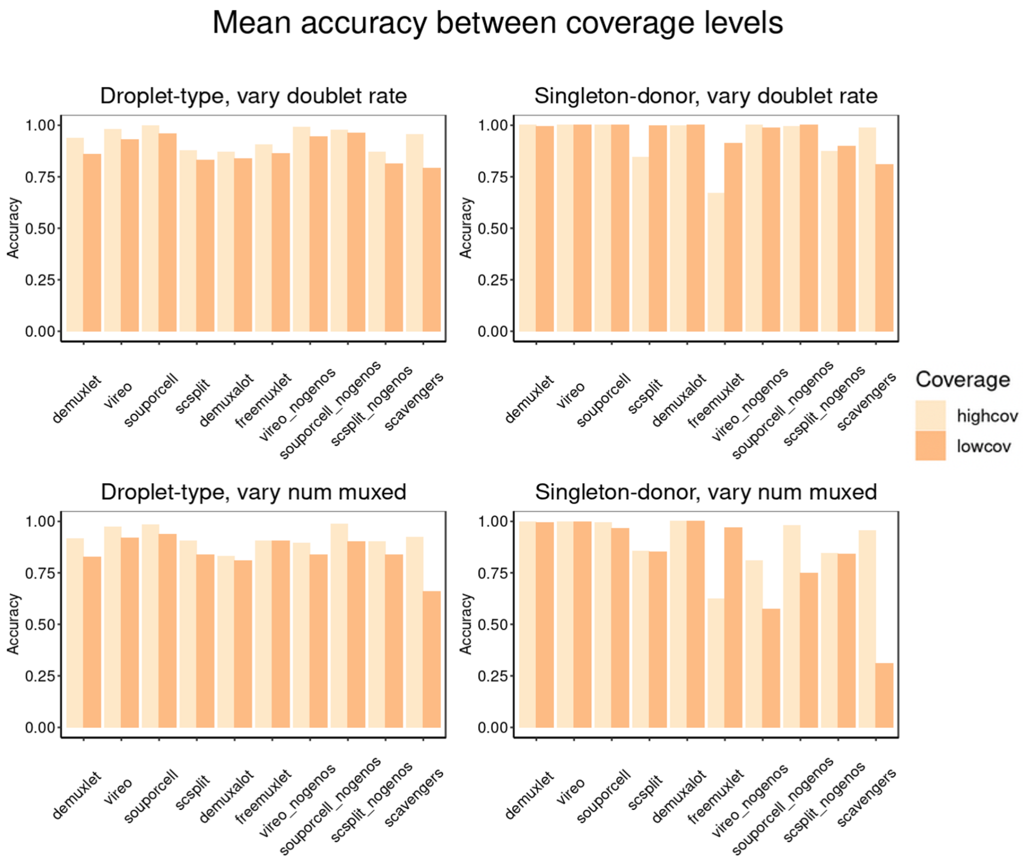


**Figure S11:** Difference between mean accuracy estimates across modalities.

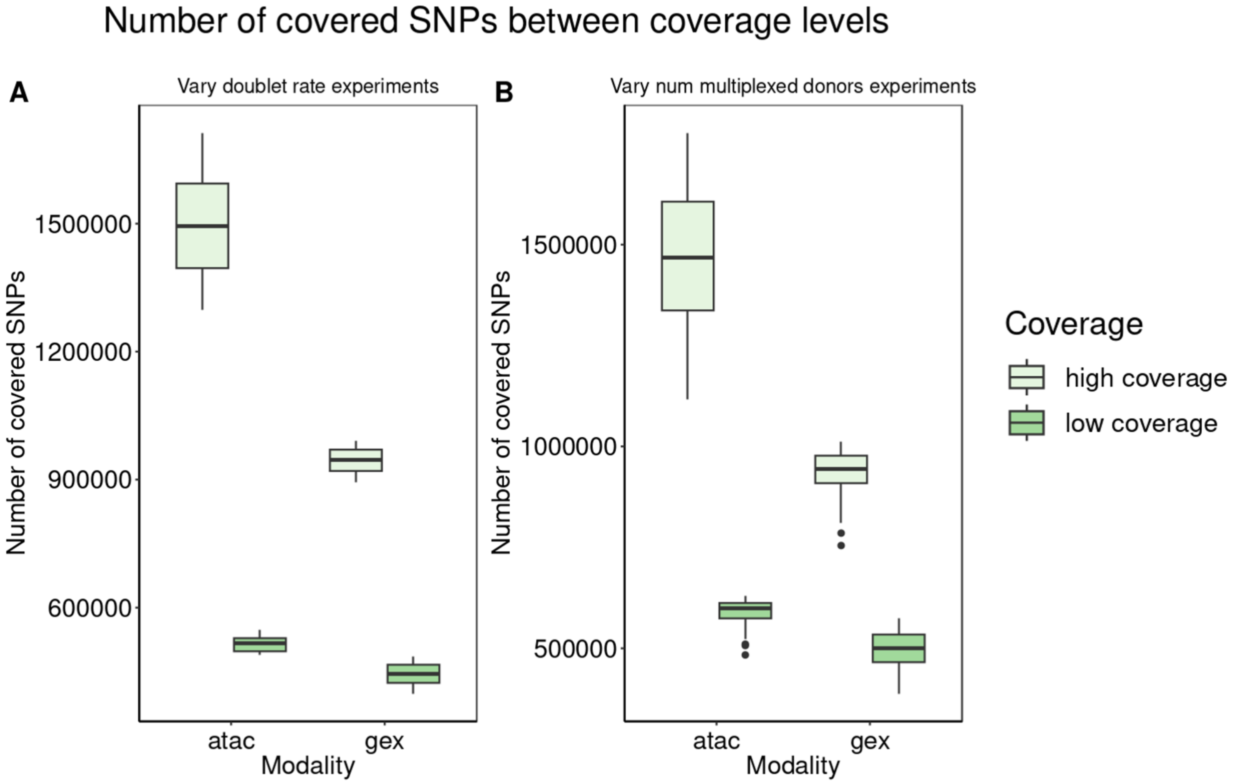


**Figure S12:** Number of SNPs covered in both modalities between high and low-coverage settings.

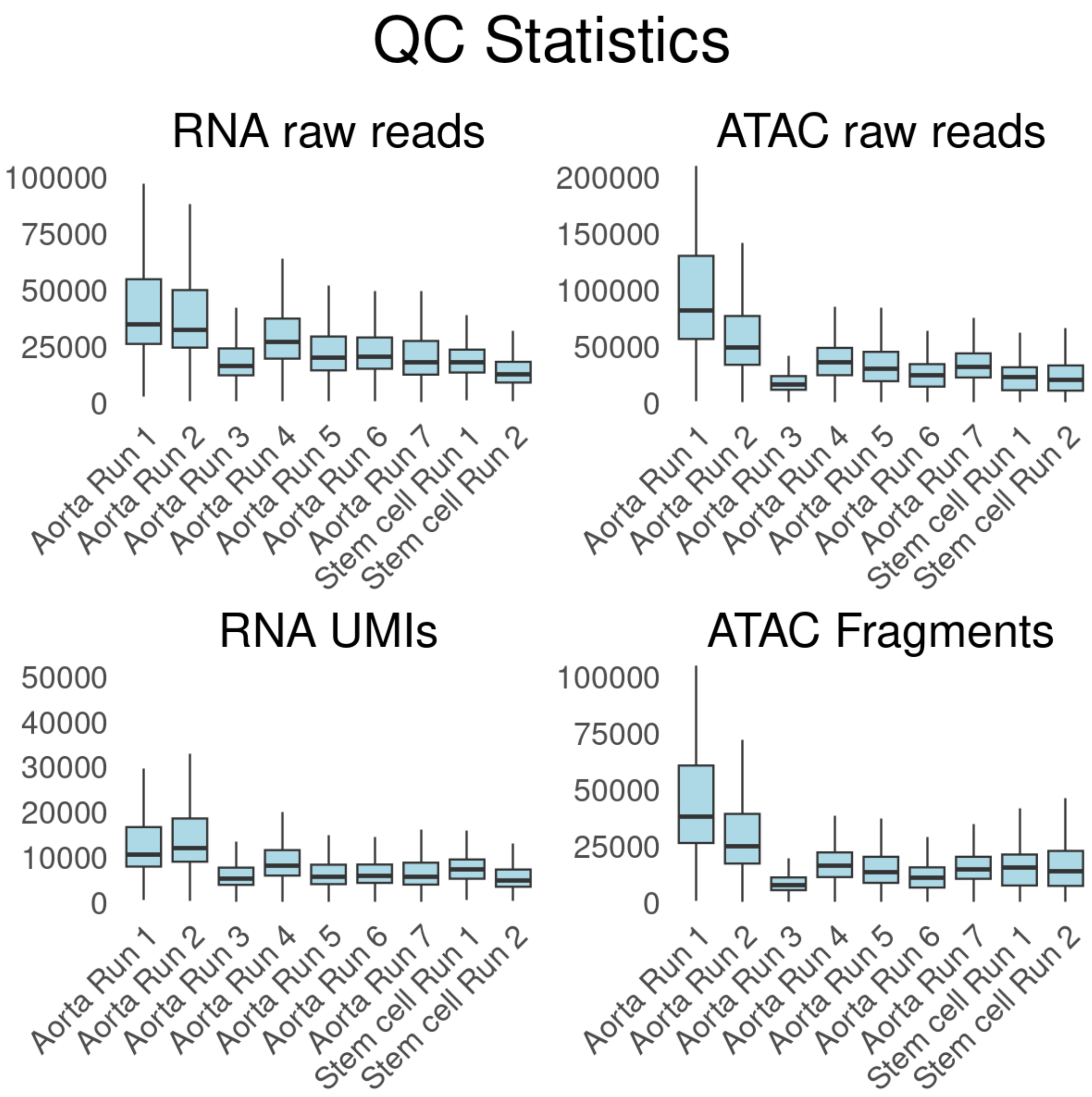


**Figure S13:** QC metrics for both real datasets.

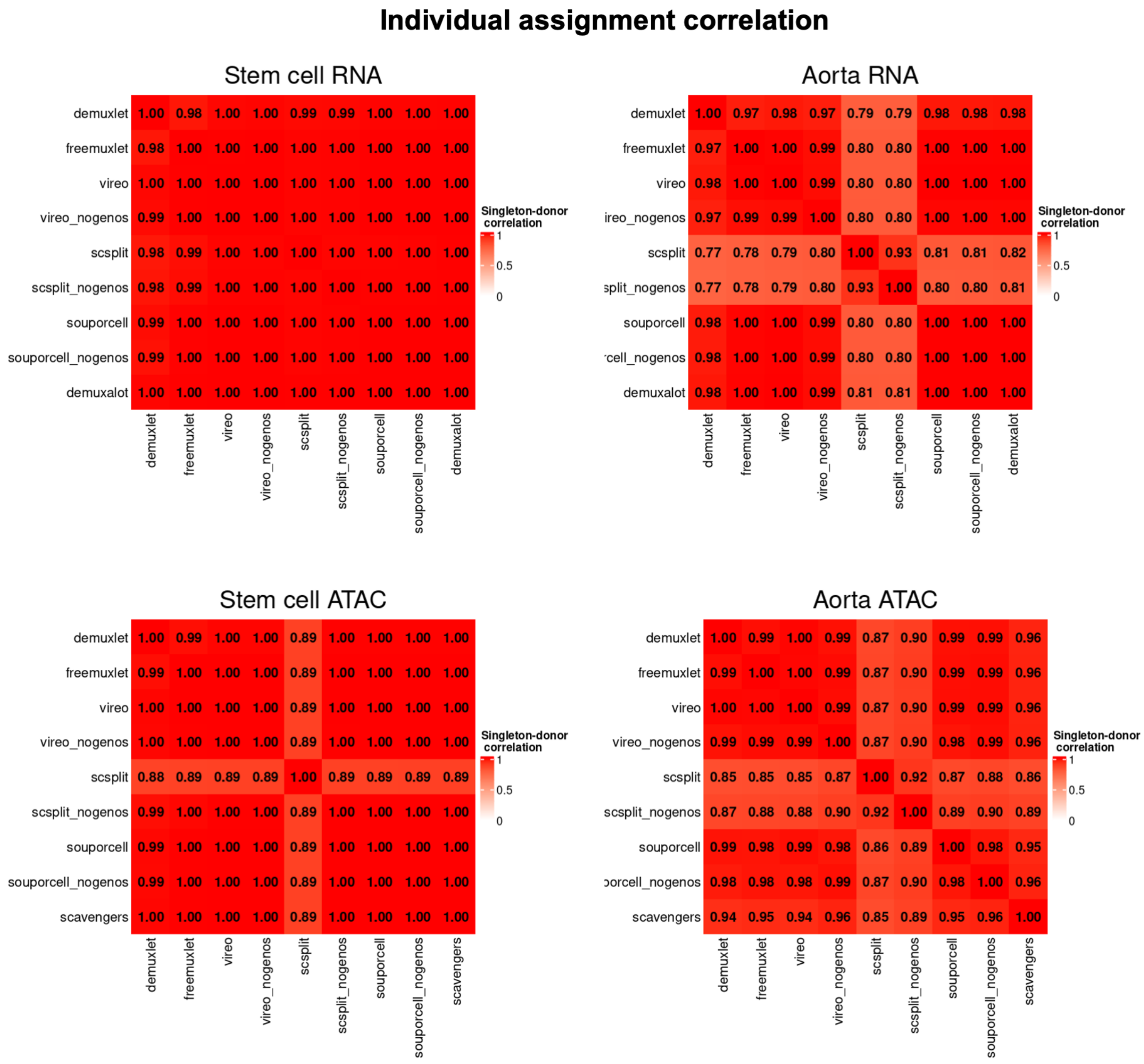


**Figure S14:** Individual assignment correlation between methods.

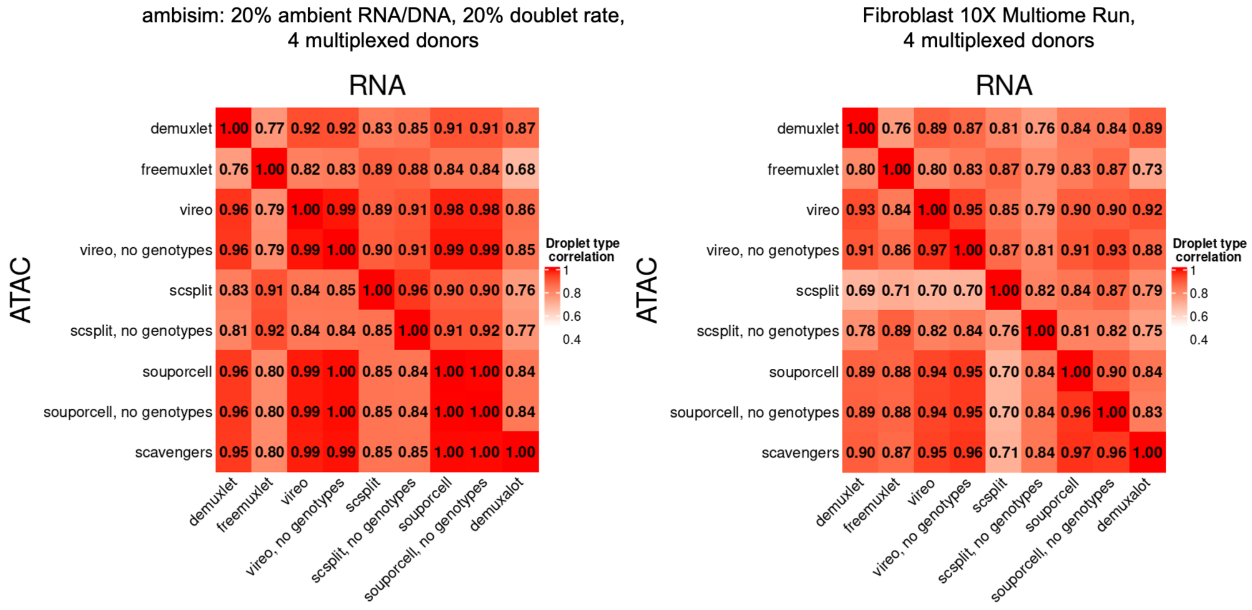


**Figure S15:** Comparison between an experiment generated by ambisim and real data.

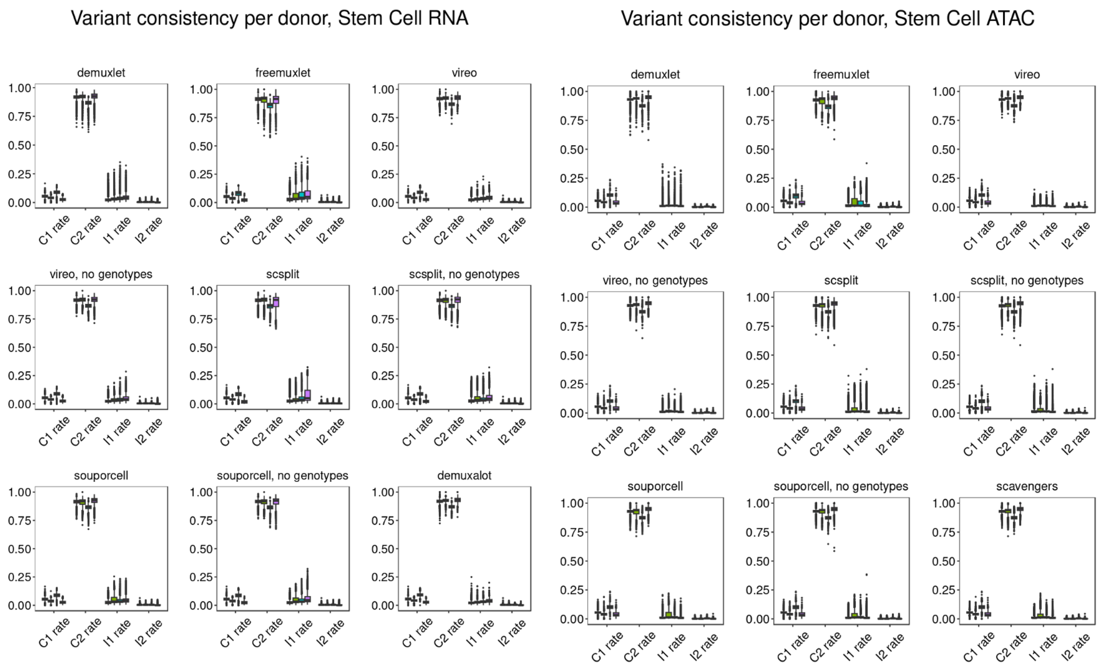


**Figure S16:** Per-donor consistency counts, stem cell dataset.

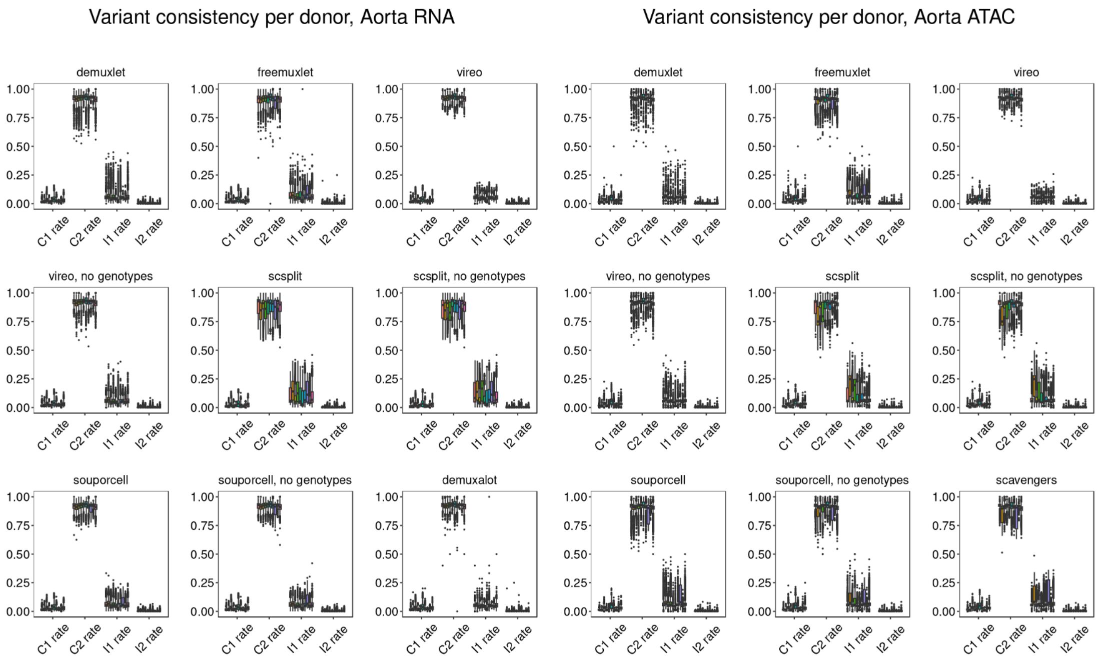


**Figure S17:** Per-donor consistency counts, aorta dataset.

### Supplemental Information

#### Comments on demultiplexing methods

We note that methods such as freemuxlet may not perform well in higher-coverage data due to cluster initialization issues and are working on adjusting parameter choices to improve the performance.

At the time of publication, several additional demultiplexing methods were not included for the following reasons:

- Mitosplitter[1]: could not be installed.
- MitoSort[2]: could not be installed.
- Dropulation[3]: demultiplexing results consistently reported >50% doublet rate. Communicating with authors to resolve issues.
- CellDemux[4]: publication date was recent.
- EnsembleX[5]: publication date was recent.

#### *ambisim*: ambient-contaminated multiplexed joint snRNA/snATAC experiments

We start by sampling cell-level attributes. First, a droplet is determined to be empty, singlet, or doublet. For each singlet, we sample a donor (or 2 donors for doublets) based on a Categorical distribution. Similarly, we sample a cell type (or 2 cell types for doublets) also from a Categorical distribution.

Then, we determine the number of RNA and ATAC reads per droplet. The number of reads per droplet is sampled from a negative binomial distribution. For empty droplets, the read coverage is sampled from a distribution mean=2 and size=0.1. For singlets and doublets, the number of reads is drawn with mean=5,000 and size=1.5 and cut within a range of 200 and 100,000.

After determining the coverage in a cellular droplet, we determine the fraction of these reads that are ambient. This fraction is sampled from a beta distribution. For the low ambient datasets, we set the shape parameters to (2, 18), resulting in a mean ambient fraction of 0.1. For the medium and high datasets, we set the parameters to (4, 16) and (6, 14) with ambient means of 0.2 and 0.3, respectively. For combined datasets, we first choose from low, medium, and high parameters randomly and then sample from that distribution.

For the ATAC modality, we also sample the number of reads which come from within peaks (intra-peak) or outside peaks (inter-peak). This fraction of reads in peaks (FRIP) is also sampled from a Beta distribution. For singlets and doublets, all FRIPs are sampled with shape parameters (4, 6). The ambient and cell reads are each split into intra-peak and inter-peak reads according to this fraction. For empty droplets, we sample the FRIP with shape parameters (1, 9).

Given the droplets donor(s), cell type(s), and RNA and ATAC coverage, we sample the individual reads. To do so, the genes and peaks are provided as genome annotations in GTF and BED format, respectively. Empty (ambient) and cell type-specific Multinomial probabilities are provided for genes and peaks. We set the distribution of the ambient profile as an average of the cell type distributions weighted by the cell type frequency. Additionally, the Bernoulli probability of a spliced read is given for the empty and per cell type in the RNA modality. We set the probability of a spliced read to 0.4 and 0.6 for cellular and ambient reads, respectively. This reflects the fact that nuclear RNA will be enriched for unspliced mRNA relative to cytoplasmic or total cell RNA. Finally, donor genotypes are provided in VCF format.

In the gene expression modality, we have the following sampling procedure. A gene is sampled from a Multinomial distribution of the gene expression profile of the cell type (or ambient). Then, an isoform for that gene is selected at random with equal probability per isoform. Finally, the read is selected as spliced or unspliced by sampling a Bernoulli distribution. The single-end read sequence is sampled uniformly within the RNA sequence of the spliced or unspliced isoform sequence.

In the ATAC modality, we employ a similar procedure but account for inter-peak reads. For intra-peak reads, we sample the peak based on the droplet cell type's Multinomial distribution. For inter-peak reads, its genomic location is sampled randomly from the genome. To generate the paired-end read, we first sample an insert size from 0 to 150 uniformly. This gives the paired ATAC-seq sequence of an ambient read.

For both ATAC and RNA modalities, the read sequences are further modified as follows. First, all variants overlapping the read alignment are selected. For each variant in a cellular read, an allele is sampled from a Bernoulli using the allele frequency of the read's donor. If the read is empty/ambient, a donor is first sampled according to a Categorical distribution and then an allele is sampled from this donor. Then, the base at the read site is replaced with this allele. To obtain the final read sequence, we sample sequencing errors with a probability of 0.01 for each base pair. Given an error, the base is replaced randomly with any of the four nucleotides.

For all simulations, we set the cell types and gene and peak probability distributions from the stem-cell multiome experiment. The top 9,000 barcodes ranked by gene expression UMIs were clustered using Seurat with a resolution of 0.5 and otherwise default parameters. This resulted in 9 cell types, and the proportion, gene expression, and peak probabilities for each were calculated empirically.
